# Euthecosomata (Mollusca, Gastropoda, Thecosomata). Taxonomic review

**DOI:** 10.1101/098475

**Authors:** Jeannine Rampal

## Abstract

The Euthecosomata Meisenheimer, 1905, holoplanktonic Mollusca with coiled or straight shell were respectively classified in Limacinoidea Gray, 1847 and Cavolinioidea Gray, 1850. In a biometrical analysis (Rampal 1973) a first change had occurd in this last superfamily: the conica shell genera *Creseis* Rang, 1828, *Boasia* Dall, 1889, *Styliola* Gray, 1850 and *Hyalocylis* Fol, 1875 were gathered into the Creseidae Rampal, 1973. Therefore it was necessary to carry on this study using molecular data. Our recent cladistic and molecular analyses as well as palaeontologic data led to a systematic and phylogenetic revision of the Euthecosomata: the Limacinoidea and of the Creseidae are not monophyletic, the other straight shells Euthecosomata are monophyletic (Corse *et al.* 2013).

The Limacinoidea are invalidated; they are split into three families: Limacinidae Gray, 1847, Heliconoididae n. fam. and Thieleidae n. fam. The Creseidae Rampal, 1973 are validated but at least there are two genera *Creseis* Rang, 1828 and *Boasia* Dall, 1889; *Styliola* and *Hyalocylis* are considered *incertae sedis.* In the Cavoliniidae Gray, 1850 there are four subfamily: Cuvierininae Gray, 1850, Cliinae Jeffreys, 1869, Diacriinae n. subfam., Cavoliniinae Gray, 1850. The Creseidae Rampal, 1973 and the Cavoliniidae Gray, 1850 belong to the Cavolinioidea Gray, 1850. The species rank of most taxa is confirmed. New genera are proposed or reinstated: *Telodiacria* n. gen., *Hyalaea* de Blainville, 1821, *Boasia* Dall, 1889. The fossil *Vaginella* Daudin, 1800 is included within the Cuvierininae Gray, 1847. The spiral fossil *Altaspiratella* Korobkov, 1966 is no longer considered part of the Limacinidae Gray, 1847.

Two phylogenetic hypotheses are analysed. According to molecular analyses in COI there is the double emergency of straight shell from two coiled shell lineages; in 28S there is monophyly; this last hypothesis we have kapt is the most parsimonious but requires some reserve and new investigations (Corse *et al.* 2013).

## INTRODUCTION

Between the Thanetian (Latest Paleocene) (Watelet & Lefèvre 1885) when the Euthecosomata first appeared and the Pliocene two morphotypes, coiled and straight-shaped shell, have diversified. Chronology and shape of fossils records coupled with phylogenetic analyses suggest that their adaptation to a planktonic environment included several stages. First the appearance of swimming organs in the spiral shell populations followed the acquisition and optimisation of straight shell of which successively evolved from conical to cylindrical pyramidal and globular. The spiral and the straight shell species were respectively classified into the Limacinoidea Gray, 1847 and Cavolinioidea Gray, 1850. In a biometrical study (Rampal 1973), the latter was separated into two families Cavoliniidae Gray, 1850 and Creseidae Rampal, 1973. Therefore a new analysis based on molecular data was necessary. The addition of our recent molecular analyses (Corse *et al.* 2013) to descriptive and evolutionary systematic data has led to a systematic revision. The Limacinoidea Gray, 1847 are broken up; the Creseidae Rampal, 1973 are redefined with the exclusion of *Styliola* Gray, 1850 and *Hyalocylis* Fol, 1875. The Cavolinioidea Gray, 1850 are restructured.

Following these recent and new insights, it was necessary to recapitulate the current knowledge about this clade: in addition to this last cladistic and molecular study, a detailed specific analysis of the phylogenetic trees for a complementary taxonomic evaluation is presented. New taxa are described and the systematic rank of several species is validated. A discussion of two phylogenetic hypotheses is tackled: single or double emergency of the straight shell Euthecosomata. Palaeontological data give valuable perspectives on these results and were compared with molecular data to estimate divergence times and also permit to follow their temporal phenotypic variations.

## MATERIAL AND METHODS

For the molecular analyses the specimens was collected during the Circum-global TARA Oceans mission (2009 – 2012) 68 stations examined. The specimens was also collected during others missions. Caribbean waters of Mexico and Belize (Eastern Coast of the Yucatan) (January 2007): 16°14’N – 21°30’N) / 86°59 W’ – 88°22’, cruise by ECOSUR (Chetunal Mexico) in collaboration with the National Oceanic Athmospheric Administration. Caribbean Sea (Virgin Islands), Missions CRER, National Oceanic Athmospheric Administration, Ship: NOAA R/V Nancy Foster; Cruise NF 08 05 (March 2008) and Cruise NF 09 03 (April 2009): 17°25’N – 18°55’N / 62°28’W – 65°14’W. Pacific Ocean, French Polynesia, Ahe Lagoon, mission FED (October 2008): 14° 49’S – 14° 45’S / 146° 35’W – 146° 33’W). Western Mediterranean Sea, Ship: Antedon, Oceanologic Center Marseille (September 2009): 43°06’N – 05°22’E. In the text are only specifed some coordinates longitude and latitude for the positive molecular analyses specimens collected during the TARA Oceans mission.

The molecular analyses are based on two sets of data: the mitochondrial Cytochrome Oxydase I gene, COI data and data set (with and without noisy sites) and the Large Subunit of ribosomal RNA 28S molecular data and gene data set (with and without noisy sites). The support values (0 to 1) show the posterior probability in the Bayesian tree, while the values (0 to 100) are the Maximum Likelihood bootstrap values. Supplementary molecular data from genebank and the Bar Coding of Life Database are using for singular genes (Corse *et al.* 2013, Tabl. 2). We also used an integrative approach to the estimation of divergence times based on the distribution of pairwise genetic distances (Corse *et al.* 2013). The estimate of taxa emergence periods is given in million years.

For cladistic analyses are also examined specimens collected during some oceanographic missions by N.O. : Thor, (1910); Dana (1921; 1930), Président-Théodore-Tissier (1957; 1958); Shoyo-Maru (1959); Thalassa (1961; 1963; 1969; 1977); Argonaut (1965), Jean-Charcot (1966; 1979; 1981); Ariadne (1966); Magga Dan (1966-1967); Coriolis (1967; 1969); Korotneff (1970; 1971); La Coquille (1971); Marion-Dufresne (1981; 1982; 1986); Missions Cyclone VI (1967), Caride 4 (1969) (00°00’ – 05° 22’S, 169°57’ – 152°32’ W).

The cladistic analysis is based on 54 variables: shell (19), parapodia-footlobe (7), head (3), pallial system (10), gill (1), alimentary system (7), genital system (6), nervous system (1); modality: presence/absence and orderly qualitative data. The matrix is available in Plos one. org Tab. S1, S2 (Corse *et al.* 2013). This study also includes a comparative morphological analysis of the radula (Microscopes Wild M5 and M10, Scanning Electron Microscope).

The main morphological characteristics are described in a key to the species (Rampal 2011). Exhaustive synonymic lists can be found in Spoel (1967) quoted on page 64.

## RESULTS

### Systematics

Two orders (two clades according to Bouchet & Rocroi 2005) Thecosomata de Blainville, 1824 and Gymnosomata de Blainville, 1824 belong to the Mollusca, Gastropoda, Euopisthobranchia (Wägele *et al.* 2013). They were united into Pteropoda then they were not considered monophyletic orders respectively related to Cephalaspidea and Anaspidea (Boas 1886; Pelseneer 1888; Meisenheimer 1905; Morton 1958; Spoel 1967; Salvini-Plawen 1970). Therefore the term Pteropoda was declined but often used. Unlike this point of view the monophyly was also admitted (Hoffman 1939; Odhner 1939) and now it is supported by molecular analyses: new clade including Thecosomata, Gymnosomata and Anaspidea (Dayrat *et al.* 2001) or Thecosomata and Gymnosomata (Jennings *et al.* 2010), with revival of Pteropoda (Klussmann-Kolb & Dinapoli 2006).

> THECOSOMATA de Blainville, 1824

Suborder Euthecosomata Meisenheimer, 1905: calcareous spiral or straight shell; two parapodia; proboscis absent (suborder studed in this text).

Suborder Pseudothecosomata Meisenheimer, 1905. Three cases for the shell: 1. calcareous spiral shell, 2. amino-acid straight pseudoconch (which calcareous spiral larval shell), 3. without shell; parapodial disc; proboscis.

> EUTHECOSOMATA Meisenheimer, 1905
>
> (Spiral Euthecosomata)

#### PALAEONTOLOGICAL SUMMARY

The earliest species known, *Spirialis mercinensis* (Watelet & Lefèvre,1885) and *Limacina heatherae* Hodgkinson, 1992 emerged during the Upper Paleocene (Thanetian) - Lower Eocene (Ypresian) (Hodgkinson *et al.* 1992). After that, fossils appeared abundantly between the Lower and Middle Eocene and showed a high degree of diversification. Few recent species, (as well as a few rare fossils) appeared only later, at the Oligocene - Pliocene: *Heliconoides inflata* (d’Orbigny, 1835), Upper Oligocene (Chattian); *Thielea helicoides* Jeffreys, 1877, Upper Miocene (Tortonian); *Limacina bulimoides* (d'Orbigny, 1836), Pliocene (Hodgkinson *et al.* 1992; Cahuzac & Janssen 2010).

#### REMARKS

The spiral shell represents the plesiomorphic state of the Euthecosomata. The caracteristic left-handed spiral shell is allocated to the neotenic extension of larval features observed in benthic forms – unique feature on the post-metamorphic ontogenesis (Bandel *et al.* 1984; Bandel & Hemleben 1995). From an anatomical perspective, the acquisition of parapodia, propelling organs derived from the foot, constitutes the first significant diversification of these planktonic Mollusca. The spiral Euthecosomata taxa were previously belonged to the Limacinoidea Gray, 1847. This superfamily was invalidated by our cladistic and molecular analyses (Corse *et al.* 2013).

#### CLADISTIC AND MOLECULAR ANALYSES

The monophyly of the Limacinoidea Gray, 1847 is supported neither by the cladistic nor by the molecular data (Corse *et al.* 2013).

Cladistic: no monophyly of the Limacinoidea where the genus *Thielea* Strebel, 1908 (characterised by a highly morphological and anatomical singularity) is the sister group to all the other Euthecosomata. The genus *Limacina* Bosc, 1817 is not monophyletic.

COI: *Thielea helicoides* (Jeffreys, 1877) + *Heliconoides inflata* (d’Orbigny, 1835) are the sister group to the strait shell Euthecosomata and did not form a monophyletic group with *Limacina helicina* (Phipps, 1774). 28S: the species of the genus *Limacina* Bosc, 1817 form a monophyletic lineage. However due to the absence of sequences *forThielea* and *Heliconoides* the monophyly of the Limacinoidea cannot be tested.

Nevertheless based on the congruence of cladogenesis and COI tree topologies and on the highly specialised morphological characteristics of *Thielea,* the Limacinoidea Gray, 1847 must be invalidated (Corse *et al.* 2013). We suggest a partition into three families: Limacinidae Gray, 1847, Heliconoididae n. fam. and Thieleidae n. fam.

> **Family LIMACINIDAE Gray, 1847**
>
> Genus *Limacina* Bosc, 1817
>
> (Figs 1; 5; 6)

**Fig. 1.**
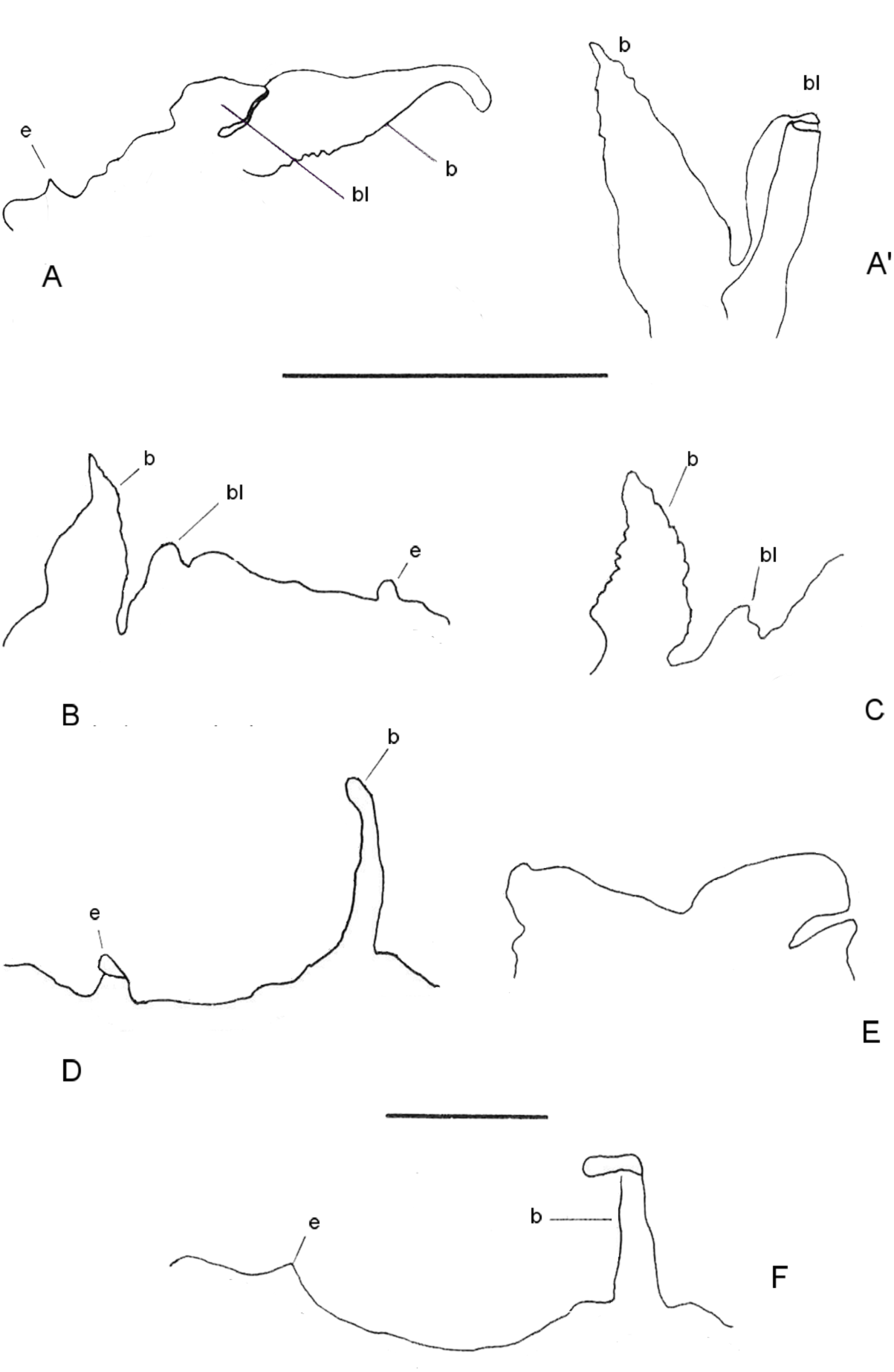
Pallial system: anterior mantle fringes and appendages (ventral view, except indication). A (dorsal view) and A’, *Limacina retroversa* (Flemming, 1823); B, *Limacina. helicina* (Phipps, 1774); C, *Limacina trochiformis* (d’Orbigny, 1836); D, *Styliola subula* (Quoy & Gaimard, 1827); E, *Creseis acicula* (Rang, 1828); F, *Hyalocylis striata* (Rang, 1828). b-bl, balancer-basal lobe; e, expansion. Scale: 0.5 mm.

Type species: *Clio helicina* (Phipps, 1774: 195)

DIAGNOSIS: Spiral shell, apex more or less hilly: apical angle = 55° – 125°, ombilic, dorsal pallial cavity, Oviparity.

*Limacina helicina* Bosc, 1817: 42.

*Spiratella limacina* de Blainville, 1824: 284 .

*Heterofusus retroversus* Flemming, 1823: 498

*Atlanta (Heliconoides) inflata* d’Orbigny, 1835:174

*Spirialis rostralis* Eydoux and Souleyet, 1840: 235

*Scaea stenogyra Philippi,* 1844:164

*Heliconoides inflatus* Hermannsen, 1846: 514

*Protomedea elata* Costa, 1861: 430

*Embolus rostralis* Jeffreys, 1870 : 86

*Altaspiratella elongatoidea* Aldrich, 1887: 83 (fossil genus)

*Thielea procera* Strebel, 1908: 85

*Limacina (Embolus) inflata* Johnson, 1934: 150

*Spiratella (Limacina) inflata* Rampal, 1964: 1

*Limacina (Thilea) inflata* Spoel, 1967: 50

Species: *Limacina bulimoides* (d’Orbigny, 1836), *L. helicina* (Phipps, 1774), *L. lesueurii* (d’Orbigny, 1836), *L. retroversa* (Flemming, 1823), *L. trochiformis* (d’Orbigny, 1836). The Limacinidae are the first Euthecosomata to appear.They are present from the Upper Paleocene (Watelet & Lefevre 1885).They are still abundant but their specific diversity decreased during the Tertiary era.

#### MATERIEL EXAMINED

TARA **Oceans mission (2009 – 2012): 674 *Limacina bulimoides* (34°93’S, 17°93E); 172 *L. lesueurii* (53°98’E, 16°95’S; 34°93’S, 17°93’E); 1487 *L. trochiformis* (16°14’N, 86° 59’W; 21°30’N, 88°22’W); 865 *L. helicina* and 169 *L. retroversa* (Antarctic).**

Western Mediterranean Sea, Caribbean Sea, South Atlantic Ocean: 88 Limacinidae.

#### DEFINITION

The spiral shell of *Limacina* has a more or less hight apex: apical angle: 55°–125°. Superficial micro-ornaments are only visible with Scanning Electron Microscope (Fig. 6A). The peristome has thin edge.The calcareous microstructure of the teloconch is prismatic and /or crossed-lamellar (Be *et al.* 1972; Rampal 1972; 1974; 1975; Richter 1976; Curry & Rampal 1979; Glaçon *et al.* 1994). The microstructure of the apex is helicoidal (Fig. 6F). There is an umbilicus more or less narrow; there is no columella. *Limacina* has a small size: w = 1 – 3 mm, h/w = 0.3 – 2.0, except *L. helicina* that can reach a width of 6 mm. The two thin parapodia are made of a single lobe each but *L. helicina* and *L. retroversa* also display a tentacular lobe on the dorsal edge of the parapodia. There is no proboscis; there is a thin cephalic lobe, and asymmetrical rhinophores. The mouth and parapodia are positioned at a similar level, like in the other Euthecosomata. The pallial cavity is dorsal. The anterior margin of the mantle is lined by a single fringe that bears some front appendages (Rampal 1975): a balancer with a basal lobe (sometimes beak-like gutter-shaped aspect) and a small expansion (Fig. 1A – C). The pallial gland includes a single layer of parallelepipedic cells on the anterior edge-second layer in the middle gland in *L. trochiformis*- and a multi-lobed prismatic cells area (Fig. 5A). Intestine with a single loop and multilamellar jaw. The median and lateral radular teeth are clearly distinct and highly differentiated (Vayssière 1915; Spoel 1967).

#### CLADISTIC AND MOLECULAR ANALYSES

The cladistic analysis gathers *Limacina helicina* and *L. retroversa* in the same group (they share the presence of a tentacular lobe on the parapodia dorsal rim similar to straight Euthecosomata, *Creseis* Rang, 1828.

28S data *L. helicina* forms a paraphyletic assemblage with *L. lesueurii, L. trochiformis, L. bulimoides.* The various analyses are not congruent with respect to *L. helicina's* position. COI data set *L. helicina* is the sister group to the Pseudothecosomata. COI data it is the sister group to *Hyalocylis.* It is difficult to make sense of these results given the current and available data about this species.

#### REMARKS

The incongruence between gene trees concerning *L. helicina* does not allow this species to be positioned within the other species. In a similar fashion, our limited available data for *L. bulimoides, L. lesueurii* and *L. trochiformis,* along with the confusion around *L. helicina* does not facilitate any clear conclusion regarding their belonging to the Limacinidae. These results constitute a first insight into the classification of this family.

#### DISTRIBUTION

In the TARA Oceans samples *L. trochiformis, L. bulimoides, L. lesueurii are* spread across all the oceans, found in temperate to tropical zones. In the Mediterranean Sea *L. bulimoides* is rare and *L. trochiformis* is abundant in the South Western and Eastern area; *L. lesueurii* is rare and restricted to the Alboran Sea: it is a marker for the presence of the Atlantic current (Rampal 1965a; 1970a; 1975). *L. helicina* and *L. retroversa* are bipolar psychrophiletic organisms. In the Arctic and Antarctic, there is a specific differentiation for *L. helicina* (Hunt *et al.* 2010). Knowledge on the diurnal bathymetric level of the Limacinidae is only partial (Rampal 1966, 1967).

> Genus *incertae sedis*
>
> *Altaspiratella* Korobkov, 1966

*Altaspiratella* Korobkov, 1966: 74.

Type species: *Physa elongatoidea* Aldrich, 1887: 83.

This fossil genus occured together with the genus *Limacina:* Early-Middle Eocene (Hodgkinson *et al.* 1992; Cahuzac & Janssen 2010); it was classified in the Limacinidae Gray, 1847.

However a striking feature that separates it from the other members of this family is the presence of a twisted columella ending with an abapical rostrum (Hodgkinson *et al.* 1992). This observation was confirmed by Cahuzac and Janssen (2010): “The columella itself is thickened and demonstrates a distinct torsion in such a way that looking into the shell’s interior is possible by a straight adapical view”. Moreover, contrary to *Limacina,* the peristome shows a notch near the columella and the umbilicus is absent. These observations are particularly significant given that Limacinidae have an umbilicus but not a columella while the Pseudothecosomata *Peracle* Forbes, 1844 is interestingly characterised by the presence of a columella and an absence of umbilicus as *Altaspiratella.* Due to lack of available data in molecular analyses for the fossil taxa, it remains difficult to definitively establish potential phylogenetical relationships between these two genera. Nevertheless, current morphological knowledge tends to support the removal of *Altaspiratella* from the Limacinidae. This genus is *sedi.*

> **Family HELICONOIDIDAE n. fam.**
>
> Genus *Heliconoides* d’Orbigny, 1835
>
> (Figs 6)

Type species: *Atlanta (Heliconoides) inflata* d’Orbigny, 1835

DIAGNOSIS: coiled shell, apex push in the last world of the shell: apical angle = 180°, ombilic, often dorsal rib, dorsal pallial cavity, prostate, pseudoviviparity.

*Atlanta (Heliconoides) inflata* d’Orbigny, 1835:174

*Spirialis rostralis* Eydoux & Souleyet, 1840: 236

*Heliconoides inflatus* Hermannsen, 1846: 514

*Limacina inflata* Gray, 1850: 50

*Limacina scaphoidea* Gould, 1852: 485

*Spirialis inflata* Adams, 1853: 59

*Heliconoides inflata* Adams, 1858: 612

*Protomedea elata* Costa, 1861: 74

*Embolus rostralis* Jeffreys, 1870: 86

*Protomedea rostralis* Fischer, 1883: 430

*Heliconoides rostralis* Monterosato, 1884: 151

*Spiratella inflata* Hedley, 1917: 106

*Limacina (Embolus) inflata* Johnson, 1934: 150

*Spiratella (Limacina) inflata* Vives, 1966: 126

*Limacina (Thilea) inflata iSpoel,* 1967: 50

*Heliconoides inflata* Janssen, 2003: 108

Species: *Heliconoides inflata* (d'Orbigny, 1835)

This species is known to be present from the Upper Oligocene onwards

#### MATERIAL EXAMINED

TARA Oceans mission (2009 – 2012): 3432 specimens (36° 21’N, 14° 30’E; 06° 03’N, 73°89’E; 16° 95’S, 53°98’E; 13° 07’S, 47°00’E; 34°93’S, 17°93’E; 09° 66’S, 09°25’W; 34°31’S, 47°59’W; 29°73’N 79° 67’W; 34°84’N 74° 13°42’S, 96° 56’ W; 03° 57’N, 154° 27’W; 12°26’N, 113°67’W; ^29°73’N, 79° 67’W; 34°84’N,74°^ 64’W).

Western Mediterranean Sea, Caribbean Sea (Yucatan; Virgin Islands), Pacific Ocean: 92 specimens.

#### DEFINITION

*Heliconoides inflata* (d’Orbigny, 1835) differs from the other spiral Euthecosomata by a planispiral shell with an involute apex pusch into the last voluminous third whorl; w = 1.2 mm, h/w= 1/3. It displays an oval projecting peristome with thin and sharp edge “peristome fortement saillant…bouche à bords très minces et tranchants” (d’Orbigny 1835: 175); t = 3.5 – 6 μm (Fig. 6 A – E). On the half of the third whorl there is a dorsal rib "large bande très hyaline terminée en croix près de l’orifice” (Vayssière 1915, Figs 153 – 155). Spoel (1967, Fig. 18A) makes clear that this structure consists of one or two oblique ribs on the middle of the last whorl prolonged by a narrow longitudinal rib “the rostrum” who ends near the pointed peristome. Sometimes is only present a rectilinear rib more or less tickened (Spoel 1967, Fig. 17 B), more or less short. Sometimes there is nothing rib: “coquille lisse” (d’Orbigny 1835: 175; 1836, Pl. 12, Figs 16 – 19). The specimens with a rectilinear rib or with two oblique and one rectilinear ribs are named type A and B (Janssen 2005). There is an opercule as the other spiral shell Euthecosomata (Rampal 1964).

The aragonitic fibres of the ribs show a chaotic structure on the external side and a more or less prismatic structure on the internal one (Fig. 6 D). The prismatic or crossed-lamellar structure of the aperture wall is similar to the rest of the teloconch (Fig. 6 B, C) but thinner. Nevertheless this area has been compared to a glassy membrane (Rang 1828) comparable with a “columellar membrane” (Spoel 1967).

The genital system is different from the rest of all the spiral shell species: firstly there is no albumine and mucus gland, no penis, but there is a prostate where the spermatophores develop, which is directly linked to the hermaphrodite gland; secondly, embryos are attached to the internal walls of the mantle (pseudoviviparity) where they develop and the offspring is laid as veliger larvae (Lalli & Wells 1973; Wells 1978).

#### MOLECULAR ANALYSES

COI: *Heliconoides inflata* and the Limacinidae are not monophyletic. It is the sister group to *Thielea helicoides;* these two species are not monophyletic with the Limacinidae; they are the sister group to the Cavolinioidea (not including *Hyalocylis* Fol, 1875, *Cavolinia* Abildgaard, 1791 and *Diacavolinia* Spoel, 1987). The relation between *Heliconoides inflata* and the other spiral species cannot be clarified. Using COI barcoding Jennings *et al.* (2010) shows that *Heliconoides inflata* is the sister group to *Creseis* Rang, 1828.

#### REMARKS

All of the present highly specific results support a separation of *Heliconoides inflata* from the other spiral Euthecosomata. The presence of two distinct lineages (1.00/100) in the Mediterranean and in the Caribbean Sea suggests the existence of two different molecular species geographically isolated. The highly specific morphological, anatomical, biological (pseudoviviparity) and molecular characteristics of *Heliconoides* prove its position into a new family. These results are corroborated by our present molecular analyses which make out four geographical species in the *inflata* group: Atlanto-Mediterranean, Indo-Pacific, South Eastern Atlantic and North Indian species entity; this last is only a little different to the Atlantic entity (one base) but COI confirm his difference (unpublished data). It is interesting to remind that *Heliconoides* is appeared from the Upper Oligocene (much later than *Limacina* which appears during the Upper Paleocene).

#### DISTRIBUTION

The species of the *inflata* group (many distinct geographical species) have a circumglobal distribution (except polar and boreal area). In the TARA Oceans samples this group is very frequent and abondant: 75% of positive prelevements, `about` 50% of the spiral Euthecosomata; maximal abondance observed in the Central Pacific Ocean.

This species have diurnal bathymetric range. Generally meso-infrapelagic, the populations have daily variations as well as seasonal migrations (linked to the laying period) during which they tend to be epipelagic (Rampal 1964; 1966; 1967; 1975: 371-379).

#### NOMENCLATURE

Janssen (2003) considers the fossil genus *Skaptotion* Curry, 1965 synonymous to the genus *Heliconoides* d’Orbigny, 1935:” I retain all forms with a apertural reinforcements in a single genus *Heliconoides’’.*

However, *Heliconoides inflata* has a thin apertural margin (see the definition above) that differs quite significantly from the apertural margin « thickened and expanded into a platform” described by Curry (1965). As a consequence according to the criterion defined by Janssen (2003), *Skaptotion* Curry, 1965 would not be part to the genus *Heliconoides.* Our molecular analyses support the position of *Heliconoides* into a new family Heliconoididae. Do to the lack available molecular analyses data for the fossil taxon it is impossible to check up their relation.

> **Family THIELEIDAE n. fam.**
>
> Genus *Thielea* Strebel, 1908
>
> (Figs 2; 5; 7)
>
> Fig. 2. Pallial system: anterior and lateral mantle fringes and appendages (ventral view, except indication). A, ( oral view schema) *Thielea helicoides* (Jeffreys, 1877); B, *Hyalaea cuspidata* Bosc, 1802; C, *Clio pyramidata* Linnaeus, 1767 ( folded asymetric fringe); D, *Cuvierina columnella* (Rang, 1827); E, *Diacria trispinosa* (de Blainville, 1821); F, *Clio polita* (Pelseneer, 1888); G, *Clio pyramidata* Linnaeus, 1767; H, *Telodiacria quadridentata* (de Blainville, 1821) n. comb. ac, calypter-like appendage; aef, aif, antero-ventral, external and internal fringes; aeff, folded antero-externe fringe; b-bl, balancer-basal lobe; eb, bilobed, expansion; la, lateral appendage; vc, vacuolized cells; vl, ventral lobe. Scale: 0.5 mm. 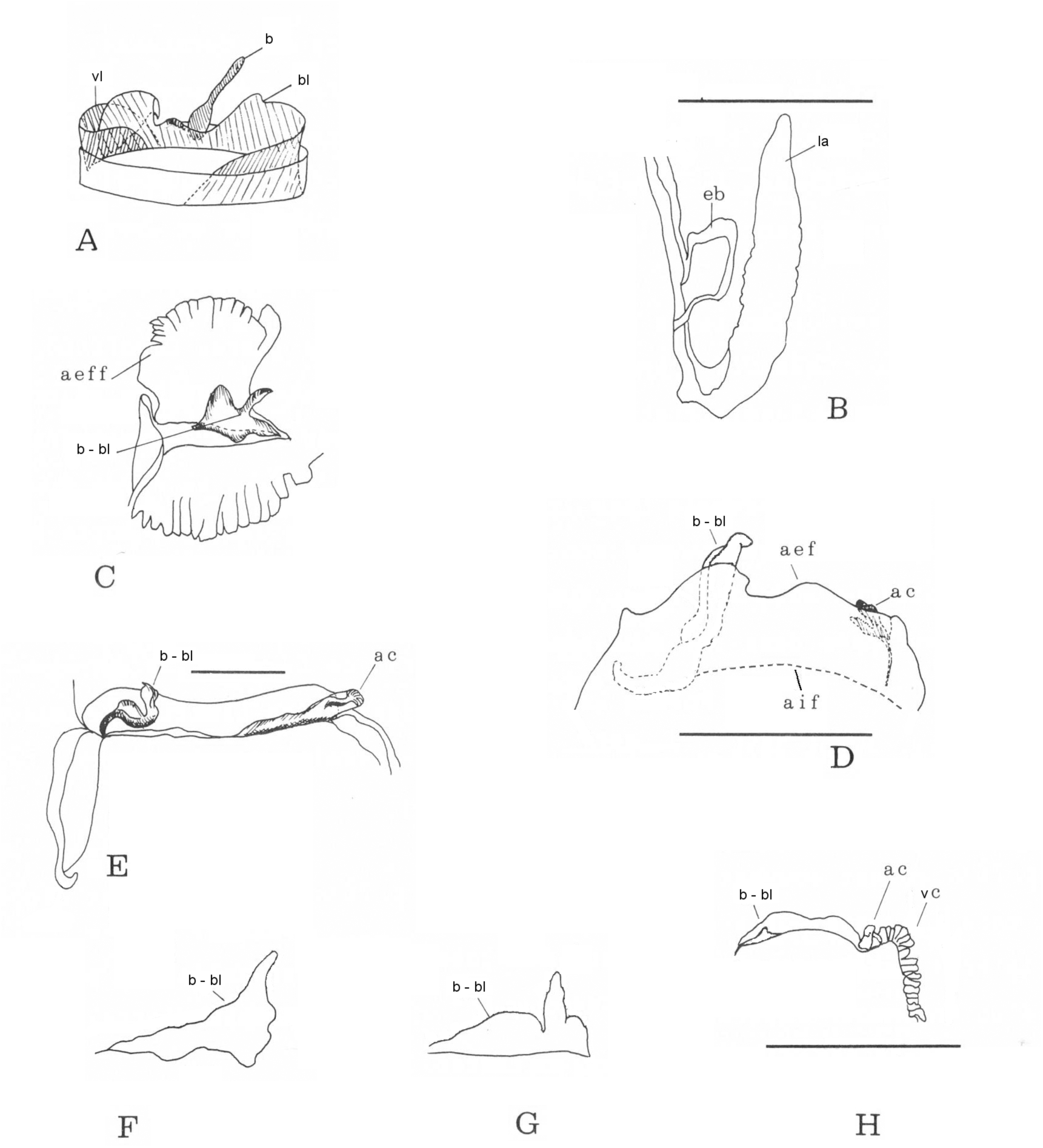

Type species: *Limacina helicoides* Jeffreys, 1877: 338.

DIAGNOSIS: calcareus spiral shell, not much apparent apex: apical angle = 125°, ombilic, cephalic lobe, lateral pallial cavity. Aplacental viviparity.

*Limacina helicoides* Jeffreys, 1877: 338.

*Thielea procera* Strebel, 1908: 85

*Thilea procera* Tesch, 1913: 22

*Spiratella helicoides* Pruvot-Fol, 1954: 117

*Limacina (Thilea) helicoides* Spoel, 1967: 48

Thilea helicoides Rampal, 1973: 1348

*Thielea helicoides* Janssen, 2004: 111

Species: *Thielea helicoides* (Jeffreys, 1877).

This species is relatively recently: Upper Miocene (Tortonian) (Janssen 2004).

#### MATERIAL EXAMINED

Data from Genbank and the Bar Coding of Life Database (Corse *et al.* 2013).

Coriolis (1967, 1969): mission Caride IV and Cyclone VI (00°00’, 169°57’E; 01°14’N, 169°49’E; 00°03N, 152°32’E).

#### DEFINITION

*Thielea* can be distinguished from the other spiral species morphologically, anatomically as well as ecologically. Size: w = 10.0 mm, h/w = 0.9. The genus has a thick shell, a very big last whorl. “The inner aperture margin is curved outwards, a trace of a rostrum is seen with a small columellar membrane like structure which is bigger in the Atlantic specimens than in the Pacific ones” (Spoel 1967). An umbilicus is present in most specimens (Spoel 1967). The soft parts are brown, the thick and folded parapodia look like a pseudo-parapodial disc and the posterior foot lobe is relatively small. There is a well developed cephalic lobe and a lateral pallial cavity. The anterior mantle border is membraneous, similar to a double fringe, where the internal fringe displays well developed and specialised anterior appendages: a balancer with a basal lobe and also a ventral lobe (Fig. 2A). These appendages were described by Bonnevie (1913) and Tesch (1946) as a "gill", similar (but not homologous) to a ctenidium, and as a "ventral body lobe" in a medio-ventral position. The pallial gland includes an anterior area with a few layers of prismatic multilobed cells and a large area of parallelepipedic cells elongated transversally except in the middle part of the gland where these cells are parallel to the antero-posterior axis (Fig. 5B). The intestine is double-looped. The median and lateral radular teeth are poorly differentiated (Vayssière 1915; Spoel 1967) and show little variety in shape and size (a notable difference from the teeth present in the other Euthecosomata); the denticles are poorly developed and irregular (Fig. 7A, A’). Another remarkable feature of *Thielea helicoides* is its mode of reproduction: aplacental viviparity (embryos are kept inside the mucus gland). This is unique among the spiral Euthecosomata (Wells 1978).

#### CLADISTIC AND MOLECULAR ANALYSES

Cladistic: *Thielea* is the sister group to all the Euthecosomata (first cladogenesis).

COI data and data set: it is not monophyletic with the Limacinidae; it is the sister group to *Heliconoides inflata* ; *Thielea* + *Heliconoides* are not monophyletic with the Limacinidae.

Together they form the sister group to the straight Euthecosomata (not including *Cavolinia* Abildgaard, 1791, *Diacavolinia* Spoel, 1987 and *Hyalocylis* Fol, 1850).

#### REMARKS

Both cladistic and COI analyses show that *Thielea helicoides* represents a separate entity within the spiral Euthecosomata and support the removal of *Thielea helicoides* from the Limacinoidea Gray, 1847. On the basis of the morphological, biological (aplacental viviparity), ecological (bathypelagic) and molecular data (distinction among the other spiral species as well as its affinities with the Cavolinioidea, apart *Creseis),* the new family Thieleidae previously proposed (Rampal 1975) is confirmed. This genus displays some plesiomorphic characteristics similar to the hypothetical benthic ancestor; it would suggest an evolutionary reversion (Corse *et al.* 2013). An analysis using an integrative approach to the estimation of divergence times would be necessary to reach a better understanding of this genus evolution.

#### DISTRIBUTION

This bathypelagic species can be found in all the latitudes of the Atlantic Ocean, as well as some rare occurrences in the South-Western Pacific Ocean.

> EUTHECOSOMATA Meisenheimer, 1905
>
> Superfamily CAVOLINIOIDEA Gray, 1850
>
> (Straight Euthecosomata)

Genera: *Creseis* Rang, 1828, *Boasia* Dall, 1889, *Styliola* Gray, 1850, *Hyalocylis* Fol, 1875, *Cuvierina* Boas, 1886, *Clio* Linné, 1767, *Hyalaea* de Blainville, 1821, *Diacria* Gray, 1847, ***Telodiacria*** n. gen., *Cavolinia* Abildgaard, 1791 and *Diacavolinia* Spoel, 1987.

This superfamily represents the second significant diversification of the Euthecosomata. The transition from a spiral to a straight shell involves deep and important morphological and anatomical changes: unwinding of the visceral mass; 180° twist of the trunk relative to the head: consecutively the pallial cavity is ventral (Boas 1886), latero-ventral in *Clio polita* and the anterior appendages are reversed. The conic shell Euthecosomata are the first to appear during the Lower Eocene.The third main diversifying event, the further optimisation into a more complex apomorphic shell occurds progressively more late.

1. All of the straight shell Euthecosomata belong to the Cavolinioidea Gray, 1850.
2. New biometrical analyses show that *Creseis, Boasia, Styliola* and *Hyalocylis* were removed from this superfamily and placed into the new family Creseidae characterised by a conical shell and some similar characteristic of the pallial complex (Rampal 1973).
3. Cladistic and molecular analyses confirm the validity of the Creseidae however limited to *Creseis* and *Boasia* (Corse *et al.* 2013); *Styliola* and *Hyalocylis* are considered genera of *incertae sedis.* Concerning the other genera of the Cavolinioidea Gray, 1850 there is incongruence: COI analyses do not support their monophyly but cladistic and 28S molecular analyses (1.00/65) show their monophyly in the Cavoliniidae Gray, 1850 and the subdivision in four subfamilies: Cuvierininae, Cliidae, Diacriinae n. subfam. and Cavoliniinae. This family represents a new radiation: outbreak from the ancestral conical teloconch to an apomorphic teloconch characterised by a dorso-ventral depression, lateral ridge then lateral slits and then a protect lip of the peristome. The times for the different lineages divergence is corroborated by the paleontology (not exactly punctual for *Clio)* (Corse *et al.* 2013).

> Superfamily CAVOLINIOIDEA Gray, 1850
>
> **Family CRESEIDAE Rampal, 1973**
>
> (Figs 1; 5)

DIAGNOSIS: straight or dorsally curved conical shell, circular transversal section, single anterior fringe on the pallial cavity, two parallelepipedic cells areas in the pallial gland.

Genera: *Creseis* Rang, 1828 and *Boasia* Dall, 1889

Genus: *Creseis* Rang, 1828: 316

Type species: *Cleodora (Creseis) virgula* Rang, 1828: 316

*Creseis conica* Eschscholtz, 1829: 17

*Criseis clava* Rang & Férussac, 1830: 261

*Hyalaea aciculata* d’Orbigny, 1836: 123

*Cleodora acicula* Deshayes & Edwards, 1836: 433

*Cresis acus* Gray, 1847: 203

*Styliola recta* Gray, 1850: 18

*Clio (Creseis) conica* Pelseneer, 1888: 50

*Clio conica* Sykes, 1905: 329

Species: *Creseis acicula* (Rang, 1828), *C. virgula* (Rang, 1828), C. *conica* Eschscholtz, 1829,

Genus: *Boasia* Dall, 1889

Type species: *Cleodora chierchiae* Boas, 1886: 202

*Clio (Creseis) chierchiae* Pelseneer, 1888: 53

*Cleodora (Boasia) Chierchiae* Dall 1889: 84

*Creseis chierchiae* Meisenheimer, 1905: 17

*Creseis (Boasia) chierchiae* Johnson, 1934:151

*Creseis virgula constricta* Chen & Bé, 1964: 194: 18

*Creseis chierchiae* Spoel, 1967: 62

*Creseis bulgia* Sakthivel, 1974: 619Species: *Boasia chierchiae* (Boas, 1886) n. comb.

#### MATERIAL EXAMINED

TARA Oceans mission (2009 – 2012): 924 Creseidae. 342 *Creseis acicula* (16°95’S, 53°98’E; 14°59’N, 69°98’E); 292 *C. conica* (35°45’N, 14°15’E); 154 *C. virgula* (73°89’N, 06°03’E); 136 *Boasia chierchiae* n. comb. (14°59’N, 69°98’E; 06°03’N, 73°89’E).

Western Mediterranean Sea, Caribbean Sea (Yucatan; Virgin Islands), Pacific Ocean, South Atlantic Ocean: 26 Creseidae.

#### PALAEONTOLOGICAL SUMMARY

The *Creseidae* are the first straight Euthecosomata to appear from the Early Eocene (Ypresian) – Middle Eocene (Chattian, Bartonian): *Creseis* almost concurrently to the fossils *Euchilotheca* Fischer, 1883, *Camptoceratops* Wenz, 1923 (which shows an ongoing unwinding of the shell), *Bovicornu* Meyer, 1887 and then *Cheilospicata* Hodgkinson, 1992 (Hodgkinson *et al.* 1992). *Euchilotheca* shows a few features not dissimilar to some taxa: presence of an internal septum, a platform-like peristome, an oblique peristome and parallel teloconch edges near the peristome. *Cheilospicata* has a conical teloconch and a tickened platform-like peristome reminiscent of the one seen in some fossil coiled shell species with thickened peristome. *Bovicornu* has a slightly coiled teloconch–a species under dispute by some authors. All these fossil genera was greatly expanded and diversified during the Tertiary era (Hodgkinson *et al.* 1992; Cahuzac & Janssen 2010).

#### DEFINITION

The teloconch of this clade represents the ancestral plesiomorphic state of the Cavolinioidea. In extant species, the conical shell is thin and more or less elongated. *Creseis:* l = 6 – 20 mm; *Boasia:* l = 2.0 – 3.5 mm. The teloconch is straight or dorsally curved, smooth or striated for *Boasia.* It has a circular transversal section, and no lateral ridge. The peristome has thin edges and its transversal section is circular. The characteristic protoconch of Creseidae is only clearly seen in *Boasia* (pear-shaped with two constrictions) and in *C. acicula* (protoconch with parallel edges and presence of thickened internally constrictions). *C. conica* and *C. virgula* have a conical protoconch more or less flared with a randomly positioned and sized constriction. The parapodia are made of a single lobe but they have a little dorsal tentacular lobe (also present in *Limacina helicina* and *L. retroversa).* The posterior footlobe is fairly small. As the other Cavolinioidea, the Creseidae have a radula with median and lateral teeth very different and a multilobed jaw (Vayssière 1915; Spoel 1967). The pallial cavity is ventral and its margin, lined by a single fringe (Rampal 1975), bears the poorly differentiated front pallial appendages (Fig. 1E). The pallial gland is composed of two areas of parallelepipedic cells (a single-layered one on the anterior margin and a multi-layered one more posteriorly); these two areas are separated by an area of prismatic cells (Fig. 5C).

#### CLADISTIC AND MOLECULAR ANALYSES

Cladistic: *Creseis* is the sister group to all the Cavoliniidae Gray, 1850

COI data (1.00/83), 28S (1.00/100): the Creseidae are monophyletic. COI data (1.00/ 83): they are the sister group to the Cavolinioidae Gray, 1850 (not including *Hyalocylis* and the Cavoliniinae Gray, 1850. 28S (0.99/78, 1.00/90): they are the sister group to the Cavoliniidae Gray, 1850.

COI (1.00/ 99, 1.00/ 100), 28S (0.95/93, 0.97/96): *Creseis conica* and *C. virgula* are the sister group. 28S (1.00/100): *C. acicula - Boasia chierchiae* n. comb. are the sister group to *Creseis conica - C. virgula.* COI (1.00/100, 1.00/99), 28S (0.97/95): *C. conica* and *C. virgula* have a specific rank. COI (1.00/91, 0.99/86): *Boasia chierchiae* n. comb. is the sister group to all the *Creseis.* 28S (1.00/96, 1.00/93): *Boasia chierchiae* n. comb. is the sister group to *Creseis acicula. Boasia chierchiae* - *Creseis acicula* are the sister group to *C. conica -C. virgula* (1.00/100).

These analyses also show distinct lineages. *Boasia chierchiae* n. comb.: COI (1.00/91, 0.99/ 86); 28S (1.00/100, 1.00/90). *Creseis acicula:* COI (1.00/99, 1.00/98); 28S (1.00/93, 1.00/99). *Creseis virgula - C. conica:* COI (1.00/91); 28S (0.95/93, 0.96/97).

#### REMARKS

These analyses support the existence of the synapomorphic Creseidae. This family represents the plesiomorphic state of the Cavolinioidea. These analyses also consolid the species level of the four species previously reinstated with biometrical analyses and characterised by the length, width, apical angle, pluri-segmented axis curve and protoconch morphology: *Creseis acicula, C. conica, C. virgula, Boasia chierchiae* n. comb. (Rampal 1985; 2002). Afterwards these taxa were also validated at the species level by Jennings *et al.* (2010), by Corse *et al.* (2013) and by Gasca & Janssen (2013). Each species is composed of several lineages found in different localities. This fact gives evidence to species groups for each species of Creseidae.

Genus: *Creseis* Rang, 1828

*Creseis conica* (Rang, 1828) and *Creseis virgula* (Rang, 1828)

No substantial regional variation was detected among Atlantic location for *Creseis conica* (Jennings *et al.* 2010). It would interesting to establish the link between the different lineages and the known phenotypes: *C. conica conica* Escholtz, 1829, *C. c. falsiparum* Rampal, 2002; *C. virgula virgula* (Rang, 1828), *C. v. frontieri* Rampal, 2002.

*Creseis acicula* (Rang, 1828).

This species displays a very stable needle-like phenotype on a world-scale basis; the specimens from diverse localities can be considered taxa of the cryptic species of the *acicula* group. These analyses show affinities between specimens from the Gulf of Aden and the South–Western Indian Ocean as well as between the ones from the Western Mediterranean and the Caribbean Sea.

Genus: *Boasia* Dall, 1889: 84

Type species: *Cleodora chierchiae* Boas,1886: 202

DIAGNOSIS: transversaly striated teloconch; pear-shaped protoconch (two constrictions). Species: *Boasia chierchiae* (Boas, 1886) n. comb.

The molecular analyses show that this species is an entity within the Creseidae. A new generic level corresponding to the hypothetical ancestor of the Creseidae is justifiable. It seems judicious to reinstate *Boasia* perfectly described by monotypy. Contrary to the other Creseidae, the teloconch of *Boasia chierchiae* n. comb. is more or less striated, the protoconch is pyriform with two extern constrictions and a round apex. It is the smallest Creseidae: *Boasia:* l = 2,0 – 3.5 mm *(Creseis:* l = 6 – 30 mm). His diploid chromosomes number is specific (Zarnik 1911, in Spoel 1967).

Specimens with smooth or almost smooth shell were described in the North Mozambic Channel (Frontier 1963; 1965). According to this author, the ringed form is the most neritic and the smooth or almost smooth specimens sometimes go beyond the continental rise; these two phenotypes reveal the polymorphism of a single species (Frontier 1963; 1965). In the Northern Indian Ocean and in the Red Sea, I have observed a large majority of specimens with only two or three striae. These phenotypes can be linked to *Creseis bulgia* (Saktivel, 1974), which is identified in the Northern Indian Ocean. Unfortunately, currently available data are not sufficient to establish the systematic level of the different *Boasia chierchiae* n. comb. phenotypes.

#### DISTRIBUTION OF THE CRESEIDAE

In the TARA Oceans samples *Creseis acicula, C. conica, C. virgula* and *Boasia chierchiae* n. comb. respectively represent 37%, 31%, 17% and 15% of the Creseidae. The *acicula* group is circumglobal, frequently found in all the oceans (40°N - 40°S) and sometimes is very abundant. *Creseis conica* is the least frequent in all the area. *Creseis virgula,* the most thermophile, is frequent and abundant in the tropical area. *Boasia chierchiae* n. comb. is a typically Indo-Pacific species. However I have observed two specimens in the Caribbean Sea: are they Panama pre-isthmatic relics?

#### NOMENCLATURE

*Creseis* acicula (Rang, 1828)

Rang (1828) describes two narrow conical shell. *Cleodora (Creseis) clava* was described as transparent shell outside water level, very elongated and sometimes irregularly flexuous, sharp apex, small and round aperture and smooth surface. l = 0,022. It is never perfectly straight (Rang 1828: 317, 318). He also describes *Cleodora (Creseis) acicula* (p.318) as a more transparent shell, needle-like, smaller than the previous one, always flexuous, with a very small aperture and smooth surface. l = 0,012. He does not think that is a young specimen of the previous one, rather a variety.

*C. clava* was the valid name used by some authors (Rang and Férussac 1830; Forbes 1844; Deshayes and Milne Edwards 1836; Cantraine 1841; Issel 1869). Souleyet (in Rang & Souleyet 1852: 56, 57), drawing from Rang's material, concluded that it is not possible to distinguish between the two species and proposes to choice the species *Creseis acicula* and the subspecies *clava*.

If there is any doubt about priority, we'll adopt the one chosen by the 1^st^ revisor (ICZN). Thus *Creseis acicula* has been used as valid by Souleyet (in Rang & Souleyet 1852) and almost universally adopted. Therefore the *C. clava* reinstatement due to the page priority (Janssen 2007) is not valid.

#### TAXONOMY

The infra-specific taxon *clava* was a source of confusions: *Creseis acicula acicula* and *C. a. clava* (Tesch 1913; Spoel 1967). *Creseis virgula* (Rang, 1828), *Creseis conica* Eschscholtz, 1829. *Criseis clava* Rang & Férussac, 1830 set apart from *acicula* (Rang, 1828) (Tesch 1948).

*C. virgula virgula, C. v. conica, C. v. clava* and *C. acicula* (Chen & Bé 1964). So the belonging to a precise species seem still unsolved.

> Superfamily CAVOLINIOIDEA Gray, 1850
>
> *Incertae sedis*
>
> Genus *Styliola* Gray, 1850
>
> (Figs 1; 5; 7)

Type species: *Cleodora subula* Quoy & Gaimard, 1827: 233.

DIAGNOSIS: conical shell, oblique longitudinal groove, circular transversal section, single anterior pallial fringe, two parallelepipedic cells area in the pallial gland.

Styliole *recta* de Blainville, 1827: 655

*Cleodora subula* Quoy & Gaimard, 1827: 233

*Cleodora (Creseis) spinifera* Rang, 1828: 313

Hyalaea *subula d’Orbigny,* 1834: 77

*Styliola subula* Gray, 1850: 17

*Clio subulata* Jeffreys, 1870: 86

Species: *Styliola subula* (Quoy & Gaimard, 1827)

This species was present from the Late Oligocene

#### MATERIAL EXAMINED

TARA Oceans mission (2009 – 2012): 173 specimens (16°95’S, 53°58’E).

Western Mediteranean Sea, Caribbean Sea (Yucatan; Virgin Islands): 73 specimens.

#### DEFINITION

*Styliola subula* has a conical teloconch (l = 7.0 – 13.0 mm) with a circular section. An oblique and longitudinal groove extends the peristome across the anterior two thirds of the dorsal side, and forms a thin pointed overhang. The peristome itself has thin edges. The pear-shaped protoconch is comparable but not similar to one observed in *Cuvierina* Boas, 1886. The parapodia are proportionally short with respect to the body size. The posterior footlobe is semi-circular like that of *Diacria* Gray, 1847 and *Clio* Linnaeus, 1767. The single fringe on the anterior margin of the mantle bears a long balancer and a short expansion (Fig. 1D). In the pallial gland the separation line between the two areas of parallelepipedic cells and the area of prismatic cells is ondulate (Fig. 5E). The radular teeth have a long and thin cuspide and their base is covered with fine denticles (Fig. 7C).

#### CLADISTIC AND MOLECULAR ANALYSES

Cladistic: *Styliola* is the sister group to *Hyalocylis* and to the Cavoliniidae.

28S mol. data. (0.83/): it is the sister group to *Hyalaea cuspidata* (Bosc, 1802) (syn. *Clio cuspidata*) and they are the sister group to the Cavoliniidae. 28S gene data set: it is the sister group to *Hyalaea cuspidata* and they are the sister group to *Cuvierina* Gray, 1847 and *Clio* Linnaeus, 1767.

#### REMARKS

These analyses all support the removal of *Styliola* from Creseidae Rampal, 1973. Based only on the comparison of the ancestral or derived features of their teloconch, affinities between *Styliola* and *Cuvierina* do not seem substantiated. Nevertheless, taking into account characteristics of the embryonic development sheds a new light on these affinities. *Styliola* could be the first rising lineage from a *Cuvierina-like* ancestor and displayed its actual morphology about 24 MY ago. Indeed, the conical plesiomorphic teloconch of *Styliola* seems a reminiscent juvenile shell with comparable protoconch of *Cuvierina. Styliola* could represent a neotenic form of *Cuvierina* (Corse *et al.* 2013). The embryonic development of *Styliola* stopped at an early stage, corresponding to the ancestral conical stage. Nevertheless the polytomy *Styliola* – *Cuvierina* do not permit an hypothetic link; so, *Styliola* would be considered *genus* of *incertae sedis.* In 28S gene data set (1.00/100) *Styliola subula* is only represented by a single species ; however these interpretations are difficult to substantiate due to the lack of results in the other analyses.

#### DISTRIBUTION

In the TARA Oceans samples *Styliola subula* represents 11% of Cavolinioidea; it is present in all the oceans (40°N-40°S) and frequently found in the South-Western and Eastern Mediterranean Sea (Rampal 1975).

> Superfamily CAVOLINIOIDEA Gray, 1850
>
> *Incertae sedis*
>
> Genus *Hyalocylis* Fol, 1875
>
> (Figs 1; 5; 7)

Type species: *Cleodora (Creseis) striata* Rang, 1828

DIAGNOSIS: striated and dorsally curved conical shell, ellipsoidal section, deciduous juvenil shell, single anterior pallial fringe, two parallelepidedic cells area, and dumbbell-shaped layer in the prismatic area.

*Cleodora (Creseis) striata Rang,* 1828: 315

*Creseis compressa Eschscholtz,* 1829: 17

*Hyalaea striata* d’Orbigny, 1836: 121

*Cleodora striata* Deshayes & Edwards, 1836: 433

*Criseis striata* Forbes, 1844: 132

*Styliola striata* Gray, 1850: 18

*Clio striata* Jeffreys, 1870: 86

*Hyalocylis striata* Fol, 1875:177

*Balantium striatum* Monterosato, 1878:115

*Hyalocylix striata* Meisenheimer, 1905:17

Species: *Hyalocylis striata* (Rang, 1828)

The genus *Hyalocylis* is known to be present from the Upper Miocene - Pliocene (Ujihara 1996). One extant and several fossils species *are* described. The fossil genus *Prehyalocylis* Korobkov, 1962 is known to be present in the Middle Eocene; it seems connected to the genus *Hyalocylis.*

#### MATERIAL EXAMINED

TARA Oceans mission (2009 – 2012) (39°53’N, 12°50’E; 35°45’N, 14°15’E): 87 specimens. Western Mediterranean Sea, Caribbean Sea (Virgin Islands), Pacific Ocean: 12 specimens.(ED mission 2008) Pacific Ocean, French Polynesia (14°49’ S, 146°35’W; 14°45’S, 33°W), surface: 3 singular specimens

#### DEFINITION

*Hyalocylis striata* has a conical teloconch dorso-ventrally depressed with a dorsal curvature, ellipsoidal transversal section, about 30 transversal striae, peristome with thin edges and no lateral ridges. l = 6.0 – 9.0 mm. The juvenile shell with conical-shaped protoconch is discarded in adults, where a blunt apex is closed by a septum, similar to *Cuvierina.* The two parapodia are very big in relation to the body size. They are rarely folded back into the shell; there is a ciliated area at their base that, again, resembles that of the *Cuvierina.* The posterior footlobe is largely reduced. The anterior margin of the mantle is lined by a single fringe as *Creseis* (Rampal 1975) and bears a long and thin balancer as well as a small triangular expansion (Fig. 1F). The pallial gland resembles that of *Creseis,* although a dumbbell-shaped layer, similar to the one in *Cuvierina,* can be found in the prismatic zone (Fig. 5D). The radular teeth have a long and thin cuspide; the trapezoidal base of the median tooth is covered with long and fine denticles (Fig.7D). In the Pacific Ocean are found singular specimens with dark marks on the edge of the parapodia (Fig.11).

#### CLADISTIC AND MOLECULAR ANALYSES

Cladistic: *Hyalocylis striata* is the sister group to the Cavolinioidea (not including *Creseis).*

COI data set: it is the sister group to all the Cavolinioidea + *Heliconoides inflata* + *Thielea helicoides.* COI data: together with *Limacina helicina,* it is the sister group to the Pseudothecosomata. 28S mol. data: it is the sister group to *Cuvierina.* 28S gene data set: it is the sister group to the Cavolinioidea (not including *Creseis).*

These molecular analyses also show that the singular specimens with dark marks on the parapodia are very likely to be undergoing speciation. (Figs 7E, E’; 11) (COI: 1.00/100; 28S: 1.00/99, 1.00/100).

#### REMARKS

The cladistic and molecular analyses support the removal of *Hyalocylis* from Creseidae Rampal, 1973, a conclusion that also was previously reached for *Styliola* (Rampal 1975: 118, 119, 122). Indeed, *Hyalocylis* only shares a few features with *Creseis,* mostly with respect to the conical shell and the pallial complex: a single fringe and only two areas of parallelepipedic cells in the pallial gland. Other traits distinguish it from Creseidae: dorso-ventral depression, presence of a dumbbell-shaped layer of prismatic cells and the shed of the juveline shell. Using 28S data, there are affinities with *Cuvierina.* To summarize, the phylogenetic position of *Hyalocylis* is difficult to ascertain and show hight instability between genes (COI, 28S) or in the same gene data (Corse *et al.* 2013). However, its ties with *Cuvierina* lineage should not be underestimated *Hyalocylis* lineage could be diverged from *Cuvierina* lineage during the Eocene and displayed its actual morphology. Despite the incongruence of results in 28S tree, the hypothesis of a common lineage *Hyalocylis-Cuvierina* does not permanently be excluded. Future studies are necessary to clarify this probleme. Nevertheless, it is deemed appropriate, for now, to consider *Hyalocylis* as genus of *incertae sedis.*

This is not the first time that the affinities of *Hyalocylis* prove difficult to assess. According to Meisenheimer (1905) “*Hyalocylix’’* originates from *Creseis* to give rise to *Cuvierina.* Bonnevie (1913) later suggested that *Clio (i.e., Clio, Creseis, Hyalocylis, and Styliola)* derives from *Clio polita* and evolves into *Cuvierina.* Klussman-Kolb and Dinapoli (2006), in their molecular analyses of the Opisthobranchia, found *Hyalocylis* basal to *Cavolinia, Diacria, Cuvierina* and *Clio.* According to Jennings *et al.* (2010) “the Euthecosomata were not monophyletic unless *Hyalocylis striata* and *Limacina helicina* sequences were excluded”.

Concerning the fossil genus *Praehyalocylis* Korobkov, 1962 (Type species; *Praehyalocylis chivensis* Korobkov & Makarova, 1962) that is included in the *Hyalocylis* lineage, it is interesting to explain their phylogenetic relationship. This fossil is known to occur from the Middle Eocene onwards (Hodgkinson *et al.* 1992). *Hyalocylis* appeared recently (6 MY) at the Upper Miocene - Pliocene (Ujihara 1996). Their relationship seems in contradiction with their morphological differences: *Praehyalocylis* shows a very big conical teloconch with no dorso-ventral depression, a circular transversal section, very thin and abundant transversal striation (about 80 striae) no dorsal curvature, a persistent juvenile shell and an elongated protoconch with a slight constriction. The integration of *Praehyalocylis* to the *Hyalocylis* lineage finds some support in our integrative approach to the estimation of divergence times reported in Corse *et al.* (2013). *Hyalocylis* lineage started comparatively early (Middle Eocene, 23MY) to the late emergence of *Hyalocylis.* It seems that *Praehyalocylis,* which appeared 17MY earlier, would represent a sister group to *Hyalocylis* with plesiomorphic ancestral caracteristics. This hypothesis thus provides an explanation for the contradiction between the belonging of *Praehyalocylis* and *Hyalocylis* to the same lineage and the gap in their respective appearance period.

#### DISTRIBUTION

In the TARA Oceans samples *Hyalocylis striata* represent 6% of the Cavolinioidea. It is often found in the inter-tropical zones of the Atlantic and Indo-Pacific Oceans. It is even more abundant and frequent in the Eastern Mediterranean Sea (Rampal 1970b; 1975). According to Jennings *et al.* (2010) “no substantial regional variation was detected among Atlantic Ocean”

> Superfamily CAVOLINIOIDEA Gray, 1850
>
> **Family CAVOLINIIDAE Gray, 1850**

DIAGNOSIS: dorso-ventral depression, triangular or elliptic transversal section, lateral ridges; double anterior fringe in the pallial cavity; three parallelepipedic cells areas in the pallial gland (transitional characterisitics for *Cuvierina).*

Genera: *Cuvierina* Boas, 1886, *Clio* Linné, 1767, *Hyalaea* de Blainville, 1821, *Diacria* Gray, 1847, ***Telodiacria*** n. gen., *Cavolinia* Abildgaard, 1791, *Diacavolinia* Spoel, 1987.

#### PALAEONTOLOGICAL SUMMARY

*Cuvierina* emerged from the Middle Eocene (Lutetian). The other genera of the Cavoliniidae Gray, 1850 emerged from the Oligocene but radiated mostly during the Miocene, long after the appearance of the earliest Cavolinioidea. *Clio* emerged during the Lower Oligocene (Rupelian) (35 MY). With regard to the other genera, they appeared between the Lower Miocene and the Pliocene: Fossil *Gamopleura* (Bellardi, 1873) (Chattian-Burdigalian); Fossil *Diacrolinia* Janssen, 1995 (Aquitanian-Langhian); *Cavolinia* (late Langhian); *Diacria* (late Tortonian) (Cahuzac & Janssen 2010). *Diacavolinia* Spoel, 1987 emerged during the Miocene – Pliocene. *Gamopleura* displays a *Cavolinia-like* shape: large and globular teloconch, noticeable lateral ridges (at times set into a groove); however, as opposed to *Cavolinia* there is no lateral slits. The peristome is ellipsoidal. *Diacrolinia* resembles *Diacria* by its extremely short juvenile shell (few millimetres) that ends in a spherical or oval protoconch and its peristome with a thick lip.

#### DEFINITION

The parapodia and the posterior footlobe more or less developed depending on the genus. There is a double fringe on the edge of the pallial cavity (Rampal 1975); the internal fringe has highly diversified anterior appendages: a balancer-basal lobe and a calypter-like appendage (Figs 2; 3). *Cavolinia* and *Diacavolinia* have lateral appendages very developed (Fig. 4). The pallial gland is composed of three areas of parallelepipedic cells (a single layered on the anterior margin and two pluri-layers connected laterally). They are separated by two areas of prismatic cells; the anterior one includes a dumbbell-shaped layer (Fig. 5F-I). Depending on the genus, the radula has a median tooth with a pointed and triangular or thin cuspide and a more or less thick and large base, which is covered on both sides with long denticles that can reach very high along the cuspide (Vayssière 1915). As the other Euthecosomata the lateral teeth are comma-shaped (Figs 8-10).

**Fig. 3.**
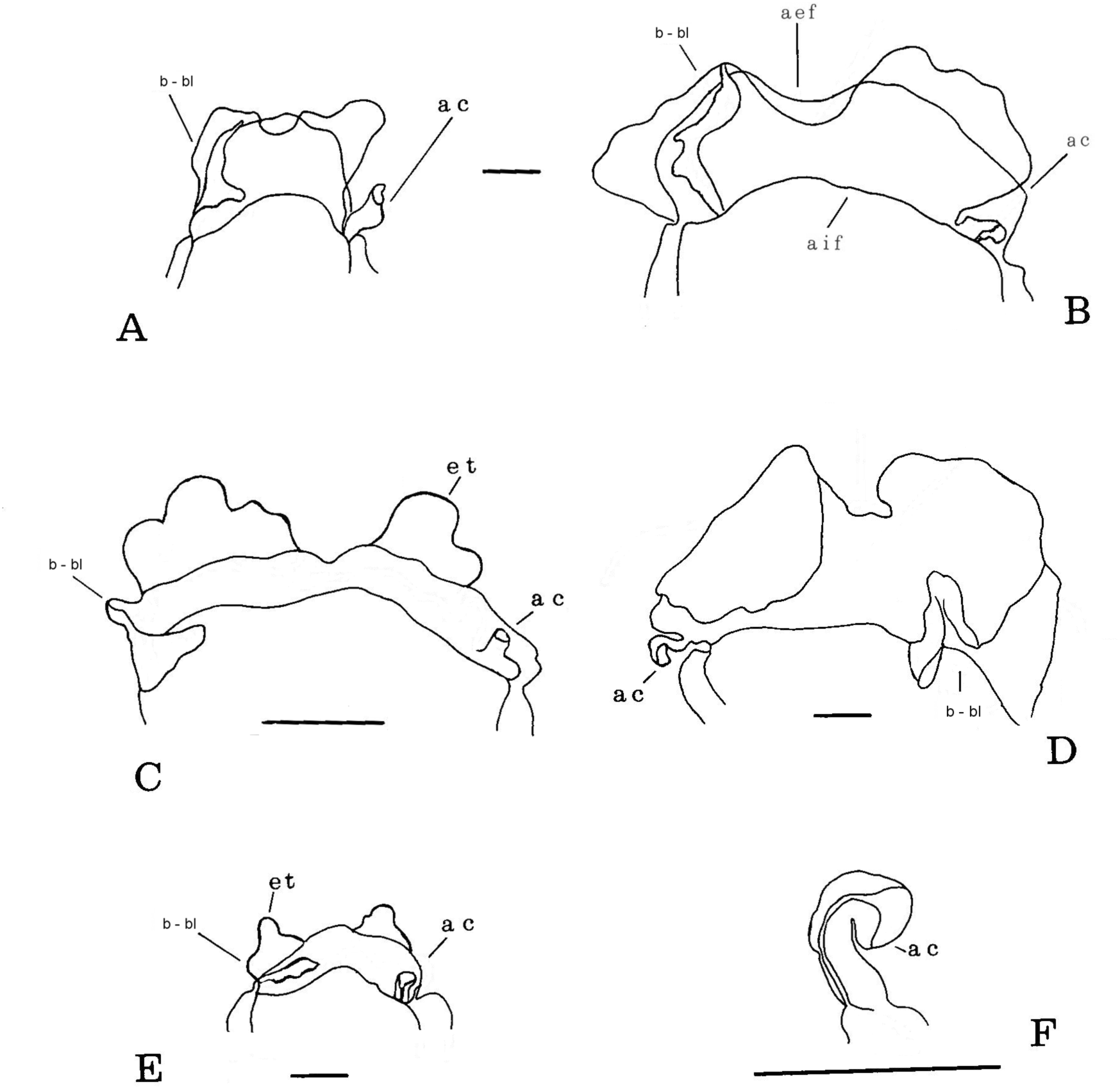
Pallial system: anterior mantle fringes and appendages (ventral view). A, *Cavolinia gibbosa* (d’Orbigny, 1836); B, *Cavolinia uncinata* (Rang, 1829); C, *Diacavolinia longirostris* (de Blainville, 1821); D, *Cavolinia* tridentata (Niebuhr, 1775); E, *Cavolinia globulosa* (Gray, 1850); F, *Cavolinia inflexa* (Lesueur, 1813). ac, calypter-like appendage; aef et aif, antero-external and internal fringes; b-bl, balancer-basal lobe; et, trilobed expansion. Scale: 0.5 mm.

**Fig. 4.**
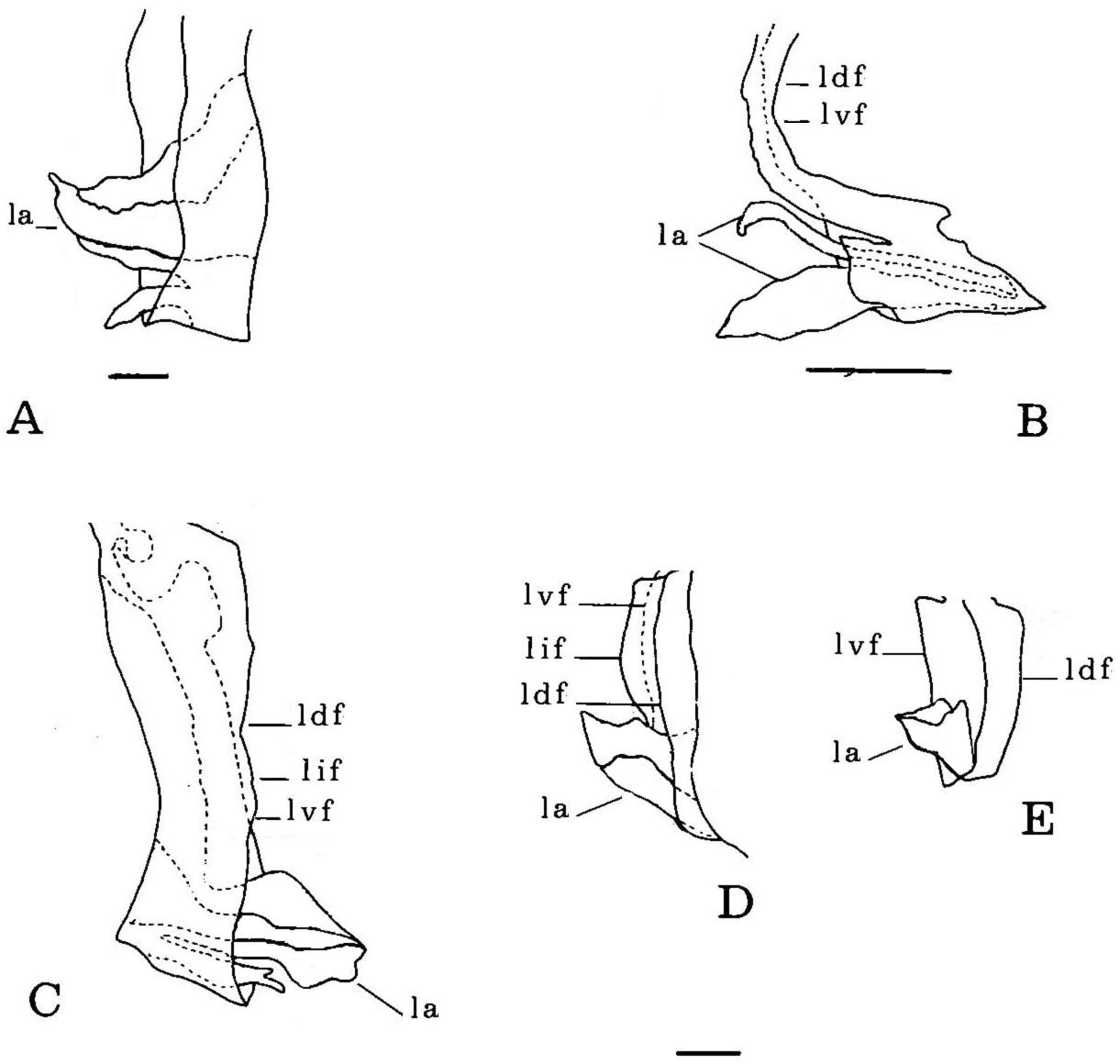
Pallial system: lateral mantle fringes and appendages (dorsal view, except indication). A, *Cavolinia gibbosa* (d’Orbigny, 1836); B, *Diacavolinia longirostris* (de Blainville, 1821); C, *Cavolinia tridentata* (Niebuhr, 1775) (ventral view); D, *Cavolinia uncinata* (Rang, 1829); E, *Cavolinia globulosa* (Gray, 1850). ldf, lif, lvfl atero-dorsal, lateral interne and latero-ventral fringe; la, lateral appendage. Scale: 0.5 mm

**Fig. 5.**
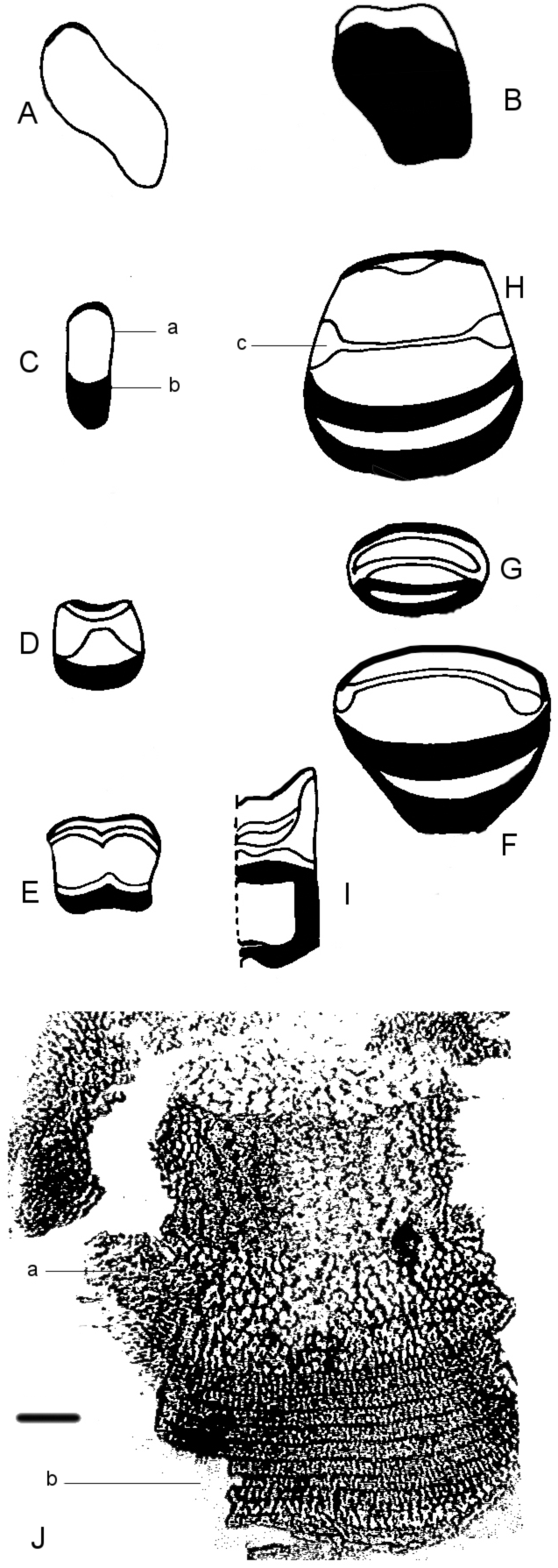
Pallial gland (A-I, schemas; dark area: parallelepipedic cells). A, *Limacina* Bosc, 1817; B, *Thielea* Strebel, 1908; C, *Creseis* Rang, 1828; D, *Hyalocylis* Fol, 1875; E, *Styliola* Gray, 1850; F, *Clio* Linnaeus, 1767; G, *Diacria* Gray 1847; H, *Cavolinia* Abildgaard, 1791; I, *Cuvierina* Boas, 1886; J, SEM microphotograph: a, prismatic cells; b, parallelepipedic cells (dark area); c, dumbbell-chaped prismatic cells area

#### REMARKS

This clade represents another significant step in the cladogenesis of the Cavolinioidea. This diversifying event characterised by an increase of the teloconch volume and a dorso-ventral depression. *Cuvierina* is the first in the radiation: transition from the ancestral conical teloconch to an apomorphic cylindrical-subcylindrical teloconch, a dorso-ventrally depression limited to the peristome, a circular transversal section, the absence of lateral ridges, a temporary juvenile conical shell with a pear-shaped protoconch (discarded during development) and the apex of the adult teloconch closed by a septum. In the other genera lengthwise lateral ridges determine the dorsal and ventral sides. *Diacria, Telodiacria, Cavolinia* and *Diacavolinia* have open lateral slits linked by closing mechanisms. Last diversification: the peristome is protected by a dorsal lip. This dorsal lip is rather thin in *Cavolinia* and thick in *Diacria* and *Telodiacria.*

The cladistic and molecular analyses infer to subdivise the Cavoliniidae Gray, 1850 into four subfamilies: Cuvierininae, Cliinae, **Diacriinae** n. subfam., Cavoliniinae.

> Superfamily CAVOLINIOIDEA Gray, 1850
>
> Family CAVOLINIIDAE Gray, 1850
>
> Subfamily CUVIERININAE (Gray, 1847)
>
> Genus *Cuvierina* Boas, 1886
>
> (Figs 2; 5; 7)

**Fig. 6.**
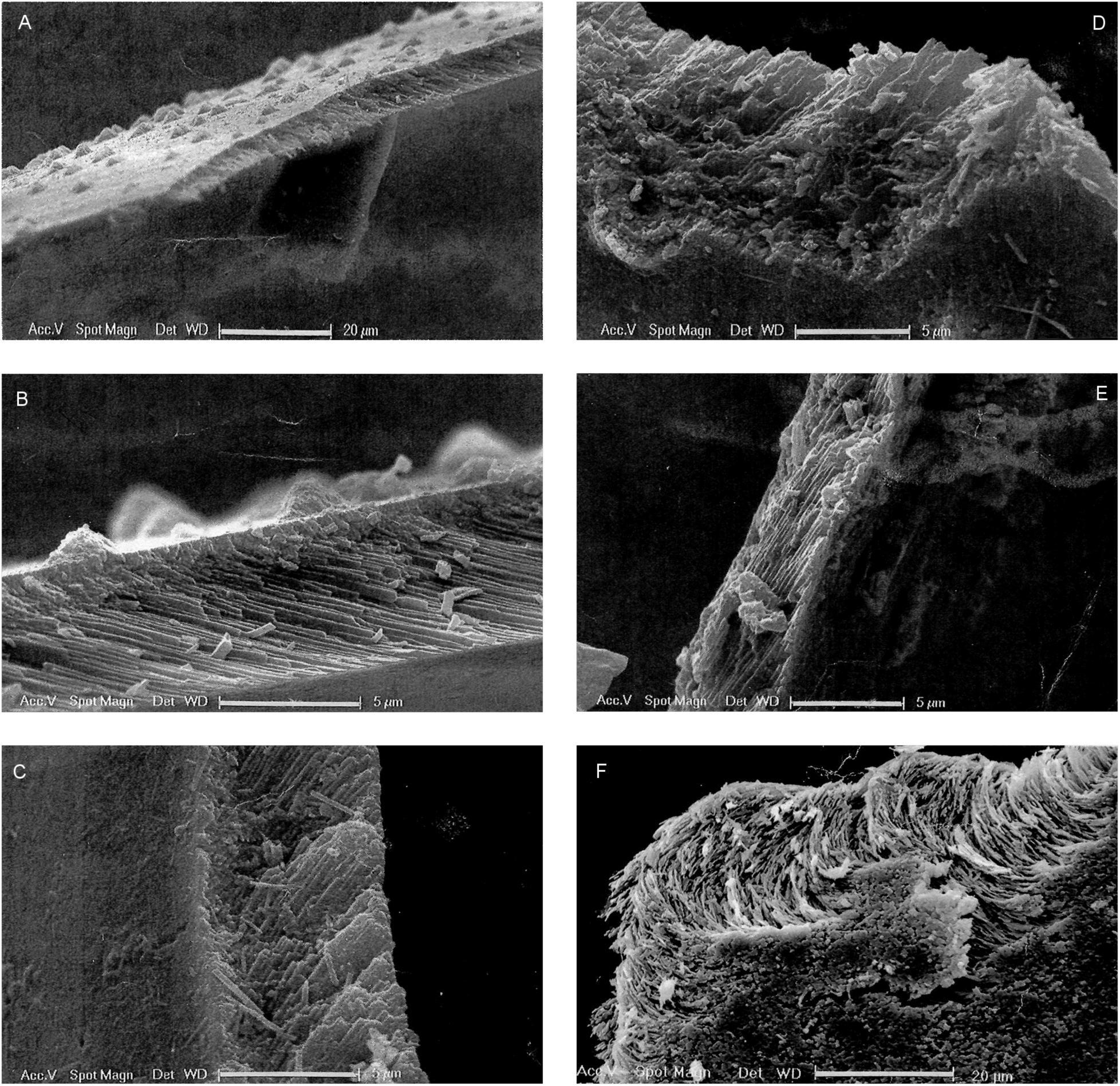
Aragonitic structure of the shell (transversal section) SEM microphotographs. A – C, *Heliconoides inflata* (d’orbigny, 1835) without rostrum: A, median point peristome; B, C, apical and ombilical margin of the peristome; D, E, *Heliconoides inflata* with rostrum: D, rostrum; E, apical margin; F, *Limacina retroversa* (Flemming, 1823), section on the apex.

**Fig. 7.**
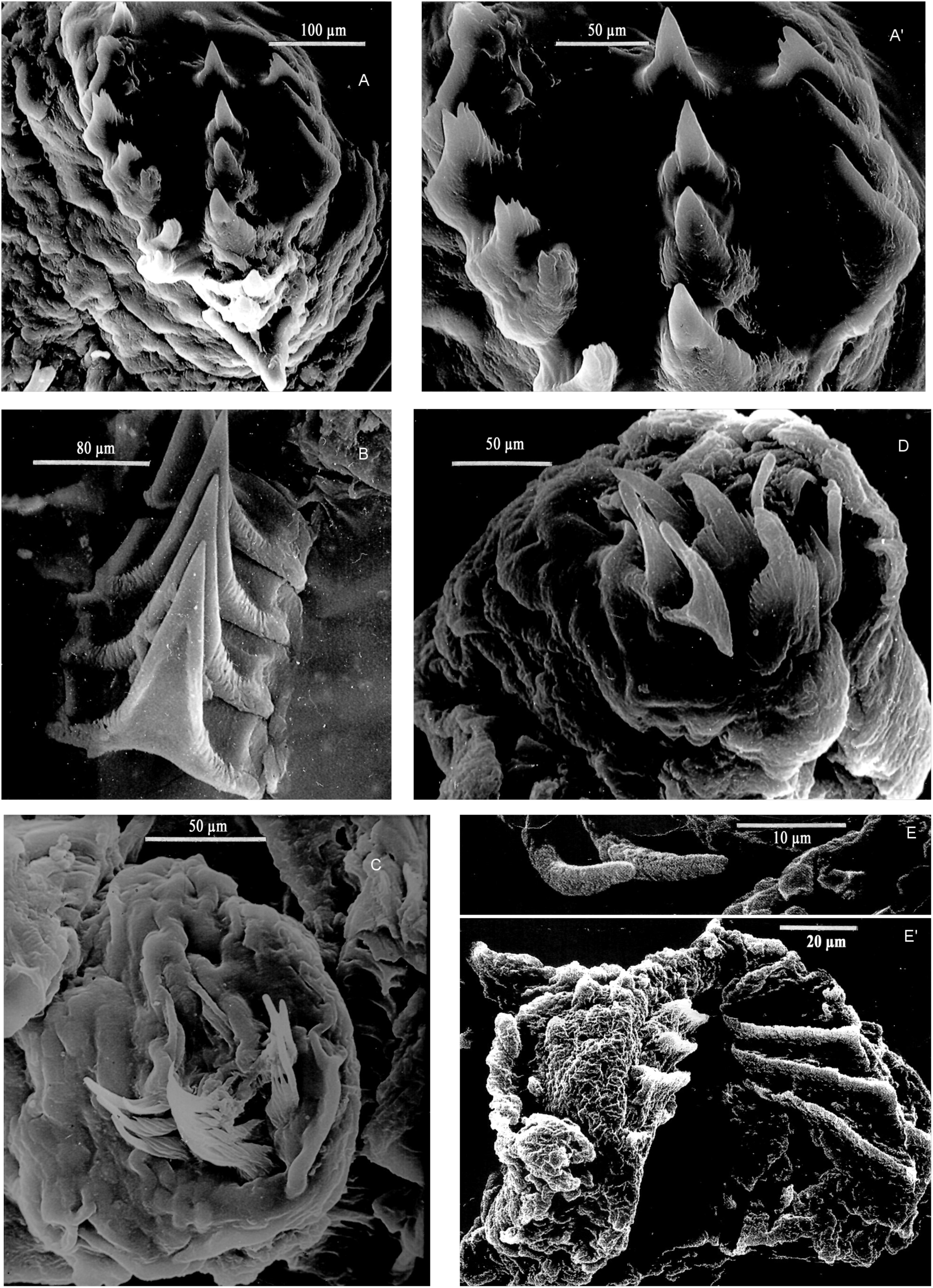
Radula. SEM microphotographs. A, A’, *Thielea helicoides* (Jeffreys, 1877); B, *Cuvierina columnella* (Rang, 1827); C, *Styliola subula* (Quoy and Gaimard, 1827); D, E, E’, *Hyalocylis striata* (Rang, 1828) (singular specimen); E, extremity of 2 teeth; E’ jaw.

Type species: *Cuvieria columnella* Rang, 1827

DIAGNOSIS: Cylindric, subcylindric or bottle-shaped teloconch, bean-shaped peristome, deciduous juvenile shell.

Species: *Cuvierina columnella* (Rang, 1827), *C. urceolaris* (Mörch, 1850), *C. atlantica* Bé *et al.,* 1972, C. *pacifica* Janssen, 2005, *C. cancapae* Janssen, 2005, *C. tsudai* Burridge *et al.,* 2016.

*Cuvierina columnella* (Rang, 1827)

Type species: *Cuvieria columnella* Rang, 1827: 322

*Cleodora (Creseis) obtusa* Rang, 1828: 317

*Cleodora columnella* Deshayes & Edwards 1836: 434

*Triptera rosea* Gray, 1847: 204

*Herse columnella* Gistel, 1848: 174

*Cuvierina columnella* Boas, 1886: 134

*Cuvierina columnella* f. *columnella* Spoel, 1970: 120, Figs. 13 C, D; 19

*Cuvierina columnella* Rampal, 2002 : 214, Fig. 1 *Cc*

*Cuvierina (Cuvierina) columnella* Janssen, 2005: 45; Figs 8-9

**Fig. 8.**
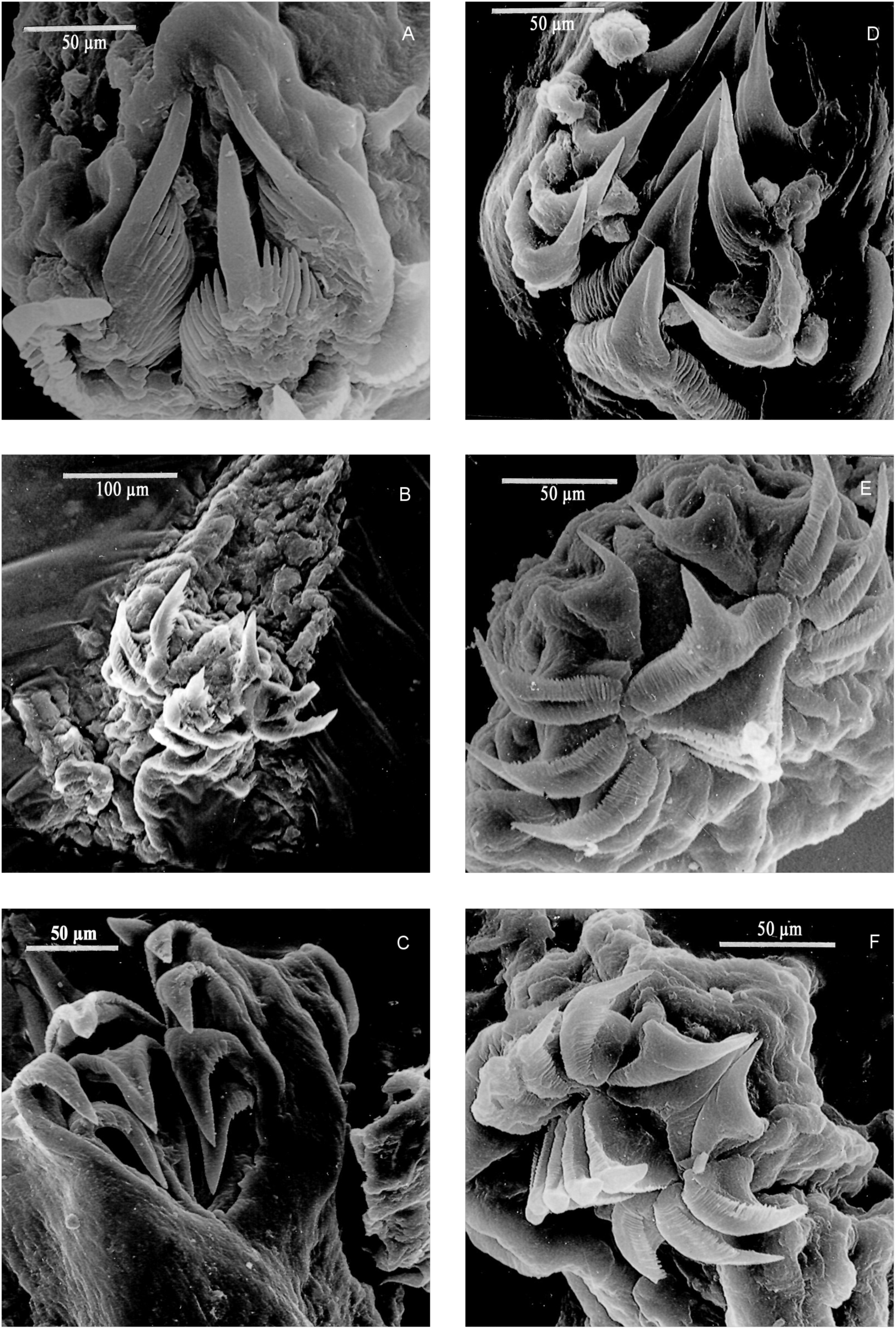
Radula. SEM microphotographs. A, *Clio pyramidata* Linnaeus, 1767; B, *Clio convexa* (Boas, 1886); C, *Clio recurva* (Childern, 1823); D, *Hyalaea cuspidata* Bosc, 1802; E, *Diacria trispinosa* (de Blainville, 1821); F, *Diacria rampali* Dupont, 1979.

**Fig. 9.**
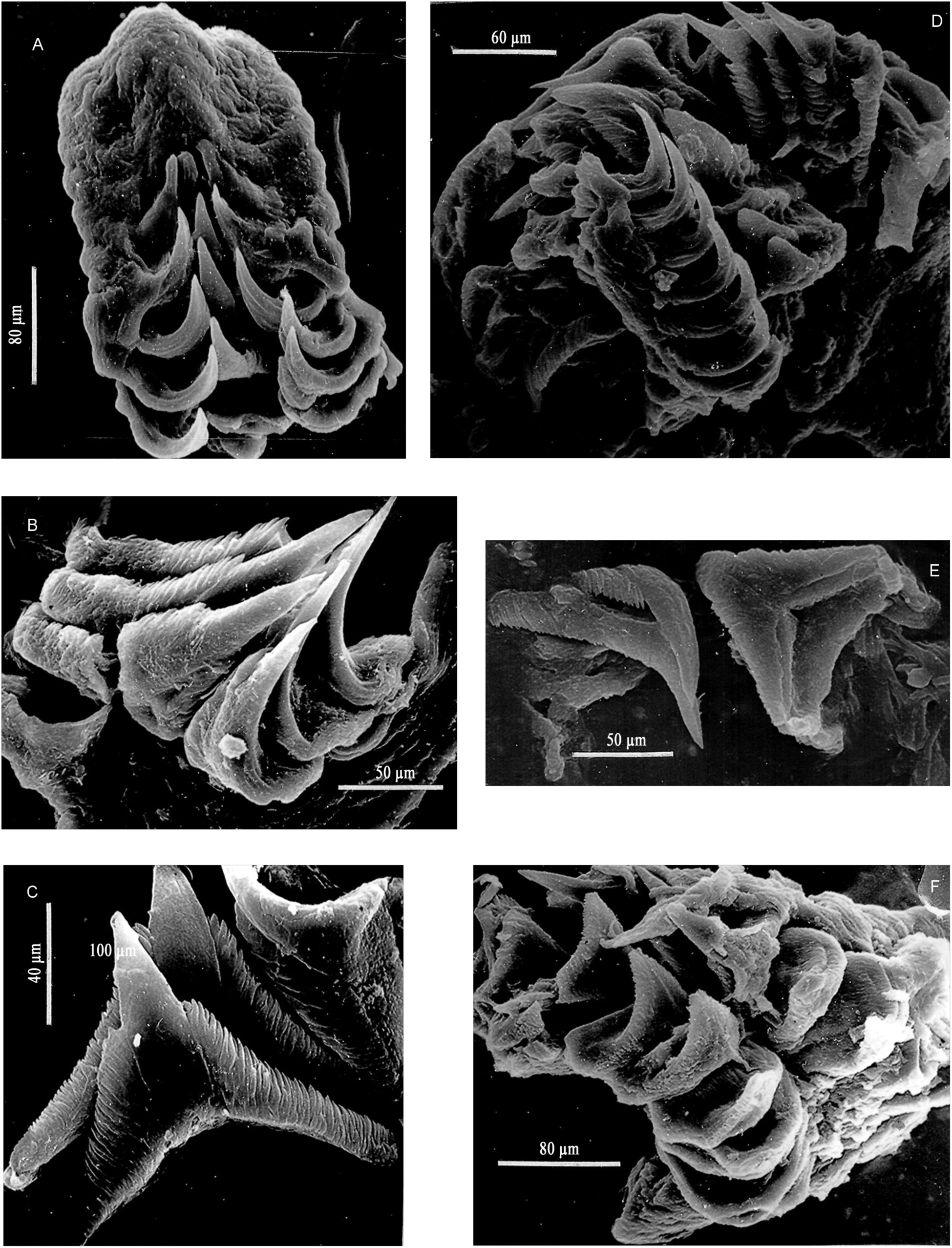
Radula. SEM microphotographs. A, B, *Cavolinia globulosa* (Gray, 1850) (A, Indian Ocean; B, Pacific Ocean); C, *Cavolinia* sp.; D, E, *Cavolinia inflexa imitans* (Lesueur, 1813); F, *Cavolinia tridentata* (Niebuhr, 1775).

*Cuvierina (Columnella) pacifica* Janssen, 2005: Fig. 14-17

*Cuvierina atlantica* Burridge, 2015, Fig. 2

*Cuvierina atlantica* Bé *et al.,* 1972

Type species: *Cuvierina columnella* f. *atlantica* Spoel, 1970: 120, Figs 13 A, B; 20)

*Cuvierina spoeli* Rampal, 2002: 214, Fig. 1Cs

*Cuvierina (Cuvierina) columnella* Janssen, 2005: 45, Figs 10-13

**Fig. 10.**
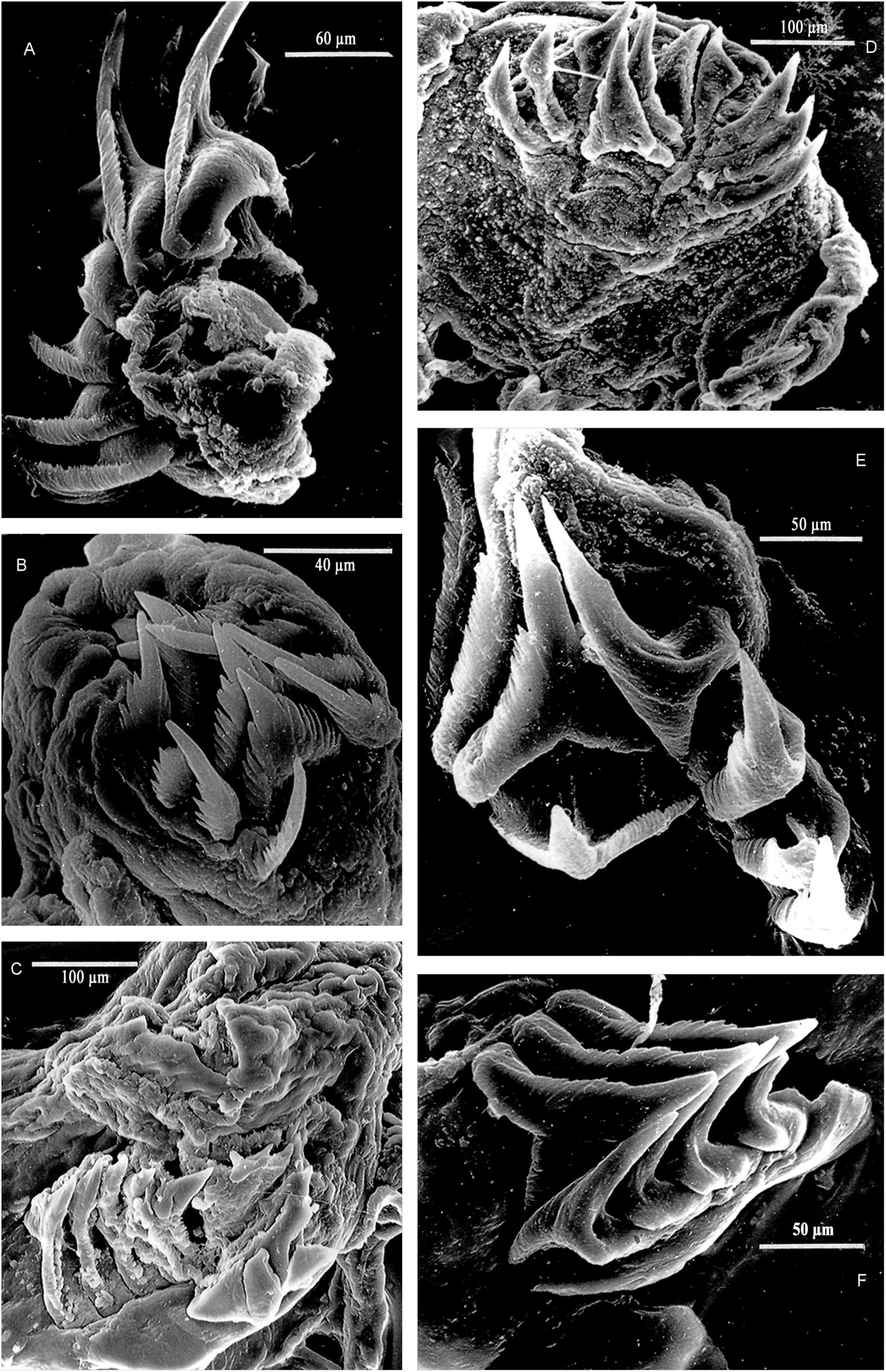
Radula. SEM microphotographs. A, *Cavolinia plana* Meisenheimer, 1905; B, *Cavolinia.gibboides* Rampal, 2002; C, *Cavolinia flava* (d’Orbigny, 1836); D, E, F, *Diacavolinia* Spoel, 1987.

**Fig. 11.**
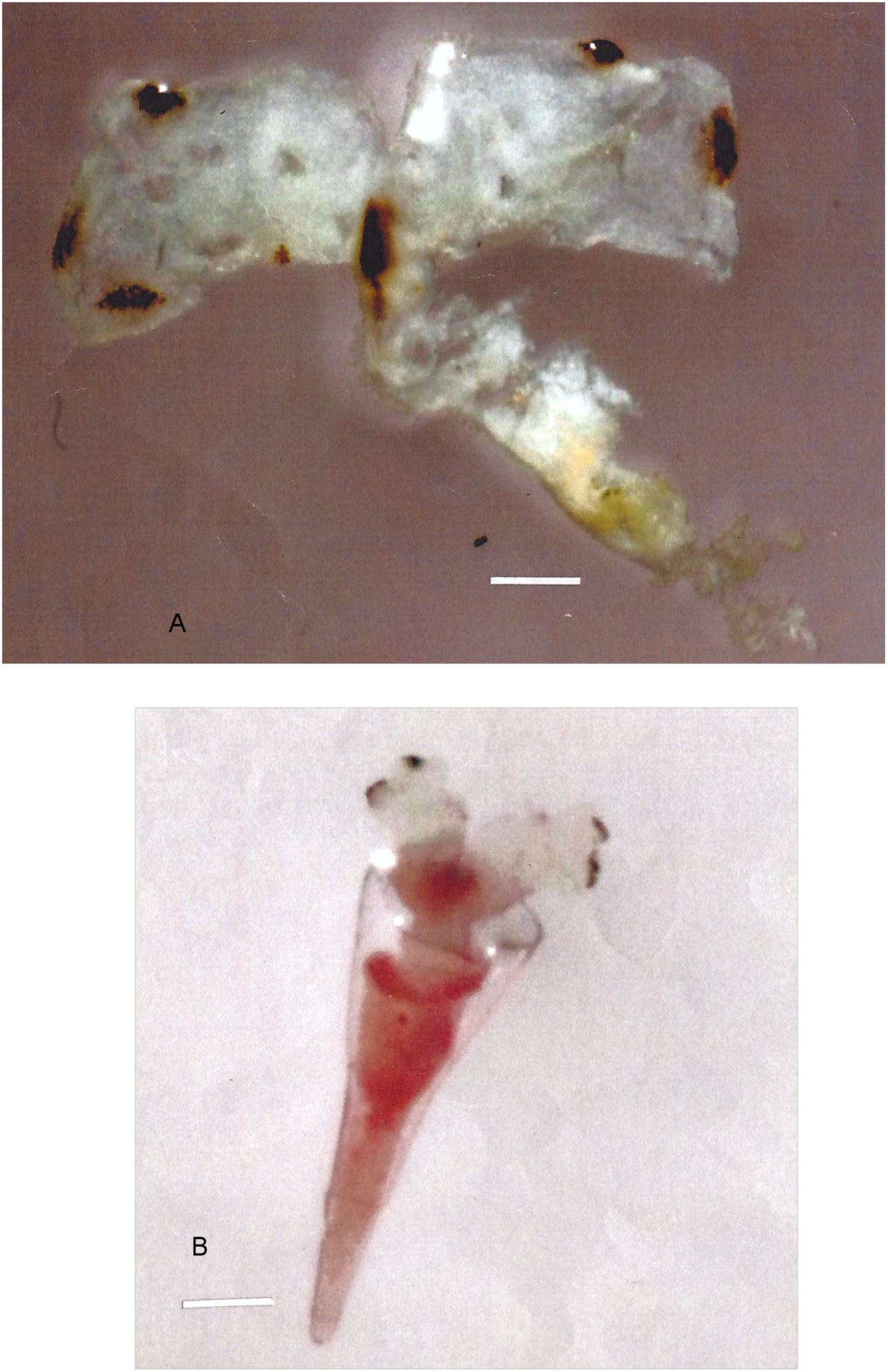
*Hyalocylis striata* (Rang, 1828) (singular specimen): A, breack adulte without shell; B, young specimen (photograph, G. Neve).

*Cuvierina columnella* Burridge *et al.,* 2015, Fig.2

*Cuvierina urceolaris* (Mörch, 1850)

Type species: *Cuvieria urceolaris* Morch, 1850: 19

*Cuvieria columnella* Souleyet, 1852: 206

*Triptera urceolaris* Adams, 1853: 55

*Cuvierina columnella* var. *urceolaris* Boas, 1886:134

*Cuvierina columnella urceolaris* Tesch, 1913: 38

*Cuvierina columnella* f. *urceolaris* Spoel, 1970 :120, Figs 13 E-G ; 21)

*Cuvierina urceolaris* Rampal, 2002: 212, Fig. 1*Cu*

*Cuvierina (Urceolaria) urceolaris* Janssen, 2005: 55, Figs 24-27

*Cuvierina urceolaris* Burridge *et al.,* 2015, Fig. 2

*Cuvierina pacifica* (Janssen, 2005)

Type *species: Cuvierina (Cuvierina) pacifica* Janssen, 2005: 46, Fig. 15

*Cuvierina columnella* Rampal, 2002: 214, Fig. 1*Cc*

*Cuvierina (Cuvierina) pacifica* Janssen, 2005, Figs 14, 15, 20

*Cuvierina pacifica* S Burridge *et al.*, 2015, Fig. 2 ?

*Cuvierina tsudai* Burridge *et al.* 2016 : 5, Fig.1 A - I

Type species: *Cuvierina columnella* Rang, 1827: 323, pl. 45

*Cuvierina columnella* Rang, 1827: 323

*Cuvierina columnella* Spoel, 1970 : 120

*Cuvierina columnella* Rampal, 2002, 214, Fig. 1*C.c.*

*Cuvierina* (Cuvierina) *pacifica* Janssen, 2005: 46, Figs 18-20

*Cuvierina pacifica* N Burridge, 2015, Fig.2

*Cuvierina cancapae* (Janssen, 2005)

Type species: *Cuvierina (Urceolaria) cancapae* Janssen, 2005 : 52 ; Fig. 21

*Cuvierina (urceolaria) cancapae* Janssen, 2005, Figs 21-23-

*Cuvierina cancapae* Burridge *et al.* 2015, Fig; 2

#### MATERIAL EXAMINED

TARA Oceans mission (2009 – 2012): *Cuvierina columnella, C. atlantica* and *C. urceolaris:* 21 specimens (16°95'S, 53°98'E; 29°50'S, 37°99'E; 26°30'S, 110°92'W).

Caribbean Sea (Yucatan; Virgin Islands): 10 specimens.

#### DEFINITION

*Cuvierina columnella* (Rang, 1827), cylindric shell, l = 7.0 – 8.0 mm; *C. atlantica* Bé *et al.,* 1972, subcylindric shell, l = 8.5 – 10.3 mm; *C. urceolaris* (Mörch, 1850) bottle shaped shell, l = 5.3 – 6.9 mm (Rampal 2002, Fig. 1, Tabl. 1); *C. pacifica* Janssen, 2005: cylindrique-subcylindric shell, l = 6,6 – 8,8 mm (Janssen 2005, Figs 14–17, 18–20); *C. tsudai* Burridge *et al.,* 2015, subcylindrique, l = 7,2 – 8,8 mm; *C. cancapae* Janssen, 2005: subcylindric-bottle shell, l=7,5 – 9,3 mm (Janssen 2005, Figs 21 – 23).

In the originally descriptions the larger Cuvierina is *C. atlantica:* l = 7,8 – 10,0 mm (Spoel 1970: 120). Paradoxally, according to Janssen (2005) and to Burridge *et al.* (2015, Fig. 2) the larger is *C.columnella,* l = 11,4 mm *(C. atlantica,* l = 8,4mm).

The peristome is bean-shaped (dorso-ventral depressed). The juvenile conical shell, circular transverse section (similar to the Creseidae teloconch) with a pear-shaped protoconch is only temporary; because it is discarded during development, the apex of the adult teloconch is closed by a septum where some remnants part of the juvenile shell can sometimes observed. The parapodia are fluffy on both sides and are almost as long as the body. The posterior footlobe is largely reduced. There is a ciliated area at the base of the parapodia (similar to *Hyalocylis).* A transitory organ is present at the base of the neck, called a cervical organ, which would have a function during copulation (Vayssière 1915). The pallial complexe is similar to the other Cavoliniidae (Figs 2D; 5I). It is strikingly different from those of the Creseidae. The median radular tooth is very different from the other Cavolinioidea; it is triangular, lined by short and tight denticles along the entire length and the base is flat and fairly thin (Fig. 7B). The nervous system of Euthecosomata is diversified with a tendency for the ganglia to merge; two remarkable features of *Cuvierina* are that the pleural ganglia can be seen externally and the existence of a second pedal commissura, as in the benthic gastropoda *Actaeon* (Boas 1886; Vayssière 1915; Hoffmann 1939).

#### CLADISTIC AND MOLECULAR ANALYSES

Cladistic: all species of the Cuvierininae are monophyletic. They are the sister group to the other Cavolinioidea (not including *Creseis, Boasia,* Styliola, *Hyalocylis*) (polytomy *Styliola-Cuvierina).*

COI data and data set (0.93/ and 0.95/): *Cuvierina* is the sister group to *Diacria* +Telodiacria + Clio. 28S mol. data (0.97/): *Cuvierina is* the sister group to *Hyalocylis.* 28S gene data set (1.00/82) *Cuvierina* is the sister group to *Clio convexa* + *C. pyramidata.* COI data and data set clearly set apart *C. columnella* and *C. urceolaris* (1.00/100). 28S data and data set (1.00/100 and 1.00/99): split between *C. columnella, C. atlantica* and *C. urceolaris.*

#### REMARKS

There is incongruence between the two 28S; consequently it is difficult to assess relationship between the other genera. Nevertheless it is clear that *Cuvierina* have a transitional role in the Cavoliniidae (affinities already shown concerning the pallial complexe) but only partial dorso-ventral depression of the peristome.Three extant species firstly considered forma level (Spoel 1970) are previously raised to a species level: *C. columnella* (Rang, 1827), *C. spoeli* Rampal, 2002 (sysn. *C. atlantica* Bé *et al.* 1972) and *C. urceolaris* (Mörch, 1850) (Rampal 2002). Our molecular analyses corroborate their taxonomic species rank (Corse *et al.* 2013). *C. cancapae* and *C. pacifica* are confirmed in molecular analyses but *C. pacifica* presents two distinct taxa: *C. pacifica* N and *C. pacifica S.* (Burridge *et al.* 2015, Fig. 2). New analyses show that *C. pacifica* N represents a distinct species *C. tsudai* Burridge *et al.,* 2016: 5, Fig. 1 A-I (Burridge *et al.* 2016).

Molecular clock analyses highlighted a basal position for the Indo-Pacific species bottle shaped *C. urceolaris.* (Corse *et al.* 2013): the split between *C. columnella* and *C. urceolaris* is relatively recent (Pliocene) and later than Janssen’s (2005) (estimation Miocene). Their distinction in two subgenera is not supported by either morphology, anatomy or the molecular clock analyses (Corse *et al* 2013). The bottle shaped fossil *C. lura* that appeared early in the Middle Eocene could incarnate the original phenotype characteristic to the Indo-Pacific Ocean.

To summarize, the genus *Cuvierina* represents the first radiation of the Cavoliniidae: transition from the ancestral conica shell to an apomorphic cylindric teloconch with a dorso-ventral depression limited at the peristome. The original morphotypes cylindric, subcylindric and bottle-shaped are conserved since the Middle Eocene.

#### DISTRIBUTION

In the TARA Oceans mission *Cuvierina* representes 6% of the Cavoliniidae; it is found between 40°N – 40°S. *C. urceolaris* can be typically found in the Indo-Western Pacific Ocean, nevertheless with rare mentions in the North-Eastern Pacific (Tesch 1948). *C. atlantica* and *C. columnella* have a widest distribution. *C. atlantica is* rather typical for the Atlantic Ocean but also may be found in the Indo-Pacific area (Spoel 1970; Rampal, 2002). *C. columnella* is the most typical for the Eastern Pacific but it also found in the Atlantic Ocean. In the Central Pacific Ocean was dradged by “La coquille” (1971) *Cuvierina columnella, C. atlantica* and *C. urceolaris* (Rampal, unpublished data).The *Cuvierina* genus is very rarely found in the Mediterranean Sea; it is restricted to the Alboran Sea where it is considered indicator for the Atlantic current (Rampal 1965a; 1970a; 1975). *C. pacifica* was found in the South Pacific Ocean and *C. cancapae* in the Atlantic Ocean (Janssen 2005).

#### PALAEONTOLOGICAL SPECIES

The Cuvierininae are occured during the Middle Eocene: *Cuvierina* Boas, 1886, *Tibiella* Meyer, 1884, *Bucanoides* Hodgkinson, 1992 (Lutetian); *Loxobidens* Hodgkinson, 1992 (late Lutetian) (Hodgkinson *et al.* 1992). *Ireneia* Janssen, 1995 (Upper Oligocene) (Cahuzac & Janssen 2010). I would add to this list the genus *Vaginella* Daudin, 1800, Lower Oligocene (Rupelian), for which possible links with the *Cuvierina* genus were previously mentioned (Rampal 1996; 2002).

- *Bucanoides, Tibiella* and *Loxobidens* have a cylindrical and cylindrical-subcylindrical teloconch with a central bulge of various sizes and have a small depression at the back of the apertural margin. There is no lateral ridge. The peristome is roughly circular or oval and is reinforced in a platform-like structure in *Loxobidens* and *Tibiella* (Hodkinson *et al* 1992). The presence/absence of this platform-like feature, which is also found in some Creseidae, is very likely to bear systematic significance, as is the case for the coiled shell species with thin/tickened peristome.
- *Vaginella* Daudin, 1800, which appeared during the Oligocene (Rupelian) - Upper Miocene (Langhian -Tortonian) was universally belonged to the Cavoliniinae. Nevertheless phenotypical and phylogenetical affinities between *Vaginella* and *Cuvierina* were already mentioned (Rampal 1996: 180; 2002: 215). Indeed, several of phenotypical features are strongly reminiscent of *Cuvierina:* a bottle-shaped teloconch with a central bulge, a dorso-ventrally depressed peristome (ellipsoïde to semi-circular) with a slightly concave ventral edge, a blunt apex closed by a septum, and a circular to sub-oval transverse section depending on the species. Several characteristics are a sign of a transition between Cuvierininae and Cavoliniinae: a tendency to a dorso-ventral depression of the teloconch (although only limited in *Vaginella)* and apparition of partial lateral ridges (along a third to half of the length in Vaginella) (dorso-ventral depression and lateral ridges run along the whole length of the teloconch in the Cavoliniinae). On the whole, this suggests a closer phylogenetical link between *Vaginella* and the Cuvierininae than with the Cavoliniinae, The comparison between *Cuvierina inflata* (Bellardi, 1873) and *Vaginella depressa* Daudin, 1800 is an illustration of this point of view. Further support for this interpretation is provided by the important synonymy between *Vaginella* and *Cuvierina.* To summarize, despite to lake of available anatomical data, this morphological data strongly suggests close affinities between *Cuvierina* and *Vaginella,* transitional form belong to the Cuvierininae.This point of view was indirectly accepted (p. 75) then declined (p. 100) by Cahuzac and Janssen (2010) and according to Janssen (2012: 405) *Vaginella* belong to the Cavoliniinae.
- *Ireneia* Janssen, 1995, Oligocene (Chattian) - Upper Miocene, displays an elliptic to semi-circular peristome reminiscent of the Cuvierininae. However, some other features set it apart: a conical teloconch, a maximum diameter at the level of the peristome and the absence of a septum (because of a persistent apex) (Janssen 1995). As a reminder, *Cuvierina* has a cylindrical or bottle-shaped shell in which the maximum diameter is never at the level of the peristome and the apex, being only temporary, is closed by a septum. This suggests that *Ireneia* that appeared fairly late in the lineage seems to represent a plesiomorphic form sister group to *Cuvierina.*
- *Cuvierina* Boas, 1886 since its emergence in the Middle Eocene (Lutetian) until the Pliocene, during which most of the species went extinct, the *Cuvierina* genus diversified significantly. The earliest occurrence, in the Middle Eocene (Lutetian), is *Cuvierina lura* Hodgkinson, 1992, showing a bottle-shaped teloconch. The one after that is *C. gutta* Hodgkinson, 1992, in the Middle Eocene (Bartonian), with a sub-cylindrical shell (Hodgkinson *et al.* 1992). The following fossil species, which appeared between the Lower and Upper Oligocene, witnessed an extensive radiation between the Lower and Middle Miocene and then significant extinction in the Pliocene (Janssen 2005). They are cylindrical, sub-cylindrical or bottle-shaped. Janssen (2005) argues that some features of *C. lura* and *C. gutta* suggest a Precuvierinidae: early emergence, absence of micro-ornaments, small size, and presence of juvenile shell remnants. This is not convincing for several reasons: these fossils appeared simultaneously with the other Cuvierininae fossils (see above), the micro-ornaments is a random characteristic (even according to Janssen), their size is within the range of the other genera dating from the Middle Eocene *(C. gutta,* l = 2.2 mm; *C. lura,* l = 1.0 mm; *Bucanoides,* l = 2.0 – 3.0 mm; *Tibiella,* l = 3.5 – 5.2 mm); as for the remnant part of juvenile shell, this can be rarely found in the other species, including extant ones. Thus, *Cuvierina gutta* and *C. lura* meet the same criteria as the other *Cuvierina.*

#### NOMENCLATURE

Spoel (1970): *Cuvierina columnella* (Rang, 1827) f. *columnella* (Rang, 1827), *C. columnella* (Rang, 1827) f. *urceolaris* (Mörch, 1850), *C. columnella* (Rang, 1827) f. *atlantica* nov. forma.

Bé *et al.* (1972): ‘Forma *columnella* is intermediate between these formae’ (Forma *atlantica* and Forma *urceolaris) ‘* in having its greatest diameter between the middle and the caudal ends of the shell…Our present study is confined to *C. columnella* forma *atlantica’* (p. 49). Afterwards (p. 58) is written *‘Cuvierina columnella atlantica* (Spoel)’: so, the taxonomically invalid *C. columnella* f. *atlantica* (ICZN) is an available subspecies, *Cuvierina columnella atlantica,* not at a new species (in accordance with the evaluation by Bé *et al.* of the three intermediate formae of *Cuvierina columnella).*

Rampal (2002: 214): biometrical analyses suggest a specific level; so, out of respect for the author of the originel description, this species was named *Cuvierina spoeli* Rampal, 2002.

Janssen (2005: 45) replaces C. *spoeli* by *Cuvierina (Cuvierina) atlantica* Bé *et al.* 1972: “according to ICZN (1999) 15.2 & 45.5.1, the name was validated as a specific group name by Bé *et al.* (1972: 58). Rampal (2002) rejected the name *atlantica* and introcuced the name *spoeli* for it. This, however, is not a replacement name, as Rampal designated another specimen as holotype, from a sample originating from 21°08’S, 55° 11’E (SW Indian Ocean). This sample, a paratype of which was available to me, belongs to *C. columnella,* which makes the name *C. spoeli to* a junior synonym of *C. columnella.* This paratype of *C. spoeli* is here designated neotype of *C. columnella* (see below)” (Janssen 2005: 45).

Janssen suggestion that *C. spoeli* is not a species does not seem grounded (apart from the use of a sample ‘available to me'). Concerning the Indian holotype locality ( South-western Indian Ocean), a border geographical zone, it is important to call back that *C. atlantica* was also found apart from the Atlantic Ocean (Rampal 2002: 214, Tabl. 2). “A geographic barrier between Indo-Pacific and Atlantic populations is unknown at present. Distribution of the two forms in the South of Africa seems to be continuous from the Indian into the Atlantic coast” (Spoel 1970: 114, 116, 118).

The molecular analyses also seem to put forward this Janssen point of view. The *C. spoeli* specific level is confirmed by our molecular analyses and the same applies *to C. columnella* and *C. urceolaris.* 28S mol. data (1.00/100); *C. urceolaris* is the sister group to *C. columnella* and *C. spoeli; C. spoeli* is the sistergroup to *C. columnella* (Corse et al. 2013, Figs 13 – 16). As a consequence the specimen named *Cuvierina spoeli* is neither a paratype nor a neotype or junior synonym of *Cuvierina columnella.* It is a synonymous of *C. atlantica.*

As to the specific group name of the genus *Cuvierina* “ a specific group takes the oldest valid species name among those of its components” (ICZN). So the specific group name of the genus *Cuvierina* is *columnella,* not *atlantica.*

> Superfamiy CAVOLINIOIDEA Gray, 1850
>
> Family CAVOLINIIDAE Gray, 1850
>
> Subfamily CLIINAE Jeffreys, 1869
>
> (Figs 2; 5; 8)

DIAGNOSIS: complet dorso-ventral depression of the triangular teloconch, lateral ridges, triangular or ellipsoidal transversal se`ction`, semi-circular posterior footlobe.

Genera: *Clio* Linnaeus, 1767, *Hyalaea* de Blainville, 1821

Genus: *Clio* Linnaeus, 1767: 1094

Type species: *Clio pyramidata* Linnaeus, 1767: 1094

*Hyaloea caudata Roissy,* 1805: 75

*Cleodora pyramidata* Peron & Lesueur, 1810 : 66

*Hyalea lanceolata* Cuvier, 1817: 381

*Hyalaea retusa* de Blainville, 1821: 81

*Clio pyramidata* Gray, 1850: 12

*Balantium australe* Gray, 1850: 15

*Clio (Clio) pyramidata* Pelseneer, 1888: 63

*Clio (Euclio) pyramidata* Bonnevie, 1913: 23

*Euclio pyramidata* Tesch, 1946: 14

*Proclio subteres* Hubendick, 1951: 3

Species: *Clio chaptalii* Gray, 1850, *Clio polita* (Pelseneer, 1888), *Clio recurva* (Childern, 1823), *C. pyramidata* Linnaeus, 1767: *C. p. lanceolata* (Lesueur, 1813), *C. p. sulcata* (Pfeffer, 1879), *C. p. martensii* (Pfeffer, 1880), *C. convexa* (Boas, 1886), *C. antarctica* Dall, 1908, *C. excisa* Spoel, 1963; *C. piatkowskii* Spoel *et al.,* 1992.

Genus: *Hyalaea* de Blainville, 1821: 9

Type species: *Hyalaea cuspidata* Bosc, 1802: 2415 (by original designation)

*Hyaloea cuspidata* Roissy, 1805: 74

*Hyalea cuspidata* Bosc, 1817: 433

*Cleodora cuspidata* Quoy & Gaimard, 1832: 384

*Clio cuspidata* Gray, 1850: 13

*Clio (Clio) cuspidata* Pelseneer, 1888: 66

*Clio (Euclio) cuspidata* Bonnevie, 1913: 22

*Euclio cuspidata* Tesch, 1946: 14

*Clio cuspidata* Spoel, 1967: 73

Species: *Hyalaea cuspidata* (Bosc, 1802) [(syn. *Clio cuspidata* (Bosc, 1802)]

#### MATERIAL EXAMINED

TARA Oceans mission (2009 – 2012): 79 Cliinae (05°99’S, 73°90’E; 16°95’S, 53°98’E; 13°08’S, 46°97’E; 39°89 N, 12°24 E).

Mediterranean Sea, Caribbean Sea (Yucatan; Virgin Islands): 15 Cliinae.

#### PALAEONTOLOGICAL SUMMARY

According palaeontological data the Cliinae appeared from the Lower Oligocene (35 MY), 37.8 MY using a pairwise genetic distance-based method to estimate divergence times by Corse *et al.* (2013). *Clio* then witnessed an important diversification at the species or infra-species levels during the Middle Miocene (Langhian). Most of the extant taxa appeared during the late Pliocene (Cahuzac & Janssen 2010).

#### DEFINITION

The Cliinae have a pyramidally shaped teloconch with a dorsal curvature of varying degree and a triangular or ellipsoidal transversal section. There are angular or flat lateral ridges but no lateral slits nor closing mechanism. In some species, the teloconch is transversally striated. The peristome is wide and its dorsal and ventral lips are not bent. There are longitudinal dorsal ribs which are absent in *Clio polita. Hyalaea cuspidata has* long dorsal and lateral spines. The protoconch is characteristic: *Hyalaea cuspidata* and *Clio polita* have a spherical protoconch (with ring in this last); the other Cliinae rather have an oval protoconch. The posterior footlobe is semi-circular. The pallial cavity is ventral; it is lateral in *Clio polita.* The fringe and the anterior mantle appendages and the pallial gland are comparable to the other Cavoliniidae but they are reduced in *Clio polita.* In the *pyramidata* group the right-hand half of the anterior external fringe of the mantle is folded (Fig. 2C). The median teeth of the radula have different forms; *C. pyramidata* have a trapezoidal-quadrangular base with numerous and long denticules and a thin cuspide (Fig. 8 A); *Hyalaea cuspidata* have a triangular base with short denticules and a triangular curved cuspide (Fig. 8D). *Clio polita* can be also distinguished from the other Cliinae by a symmetrical nervous system with fused visceral and pedal ganglia (Bonnevie 1913; Tesch 1946; Spoel 1967).

#### CLADISTIC AND MOLECULAR ANALYSES

The cladistic analysis includes: *Clio convexa, C. recurva, C. chaptalii, C. pyramidata, Hyalaea cuspidata* and *Clio polita.* Unfortunaly for the molecular analyses the taxonomic sampling and the phylogenetic trees resolution are only obtained for *Clio pyramidata, C. convexa* and *Hyalaea cuspidata*.

Cladistic: the Cliinae are monophyletic and they are the sister group to the Diacriinae n. subfam; *Clio polita* is the sister group to *Clio recurva* and to *Clio chaptalii.*

Molecular analyses. COI: the Cliinae are monophyletic; *Hyalaea cuspidata* is the sister group to the new monophyletic clade *Clio pyramidata* + *C. convexa;* the Cliinae are the sister group to *Diacria trispinosa* + *D. major.* 28S gene data set: the Cliinae are not monophyletic (in relation to *Hyalaea cuspidata); Hyalaea cuspidata* is the sister group to *Styliola subula,* they are the sister group to the Cuvierininae and to the other Cliinae; 28S mol. data: the Cliinae are not monophyletic; *Hyalaea cuspidata* is the sister group to *Styliola subula* and they are the sister group to other the Cavoliniidae (except the Creseidae). According to Jenning *et al.* (2010) in COI *Clio recurva* is geneticaly similar in the Eastern and Southern Atlantic Ocean.

#### REMARKS

In COI the Cliinae are monophyletic; in 28S they are not monophyletic in relation to the inconsistant position of *Hyalaea cuspidata.Two* species *Clio polita* and *Hyalaea cuspidata* stand out as very entity in the Cliinae.

*Clio polita* Pelseneer, 1888

The present limited data in the phylogenetic trees resolution do not permit to propose a generic break for *Clio polita.* Nevertheless this bathypelagic species presents some plesiomorphic characteristics especially the lateral pallial cavity, the reduced anterior pallial fringes, flat and blunt parapodia and the symmetrical nervous system with fused visceral and pedal ganglia. Despite the deficient molecular analyses, the morphological, anatomical and ecological characteristics suggest that this species is the witness of an ancestral plesiomorphic state of an hypothetic Cliniinae’s common ancestor.This bathypelagic species could have emerged recently from an ancient lineage and would have retained some of its features. This hypothesis could be tested using the integrative approach to the estimation of divergence times and molecular analyses.

*Hyalaea cuspidata* Bosc, 1802: 241 (Figs 2B; 5F)

The molecular analyses support a genus break for this species before named *Clio cuspidata.*

`28S: the Cliinae` are not monophyletic in relation to the inconsistante position of *Hyalaea cuspidata*. It also can be distinguished from the orher Cliinae by several features : lateral and dorsal spines of the teloconch, spherical protoconch with a terminal spine, characteristic intestin appendage, exclusive structure unique in the family (Tesch 1946; Spoel 1967); one dorsal and two bilobed lateral appendages; left lateral bilobed expansion (Rampal 1965b): calypter like appendage homologous? (Fig. 2B).

Despite his relative phenotypical stability, different sequences “were detected between regions for *Hyalaea cuspidata* (SE Atlantic vs. Southern Ocean)” (Jennings *et al.* 2010).

#### DISTRIBUTION OF THE CLIINAE

In the Tara Oceans samples the Cliinae represent 22% of the Cavoliniidae. *Clio convexa* is a characteristic Indo-Pacific Cliinae. A subspecies *Clio convexa cyphosa* Rampal, 2002 was identified in the Red Sea.

The *pyramidata* group is polytypic and polymorphic (Spoel 1963, 1967, 1969a; Rampal 1975, 2002). *Clio pyramidata lanceolata:* 62% of the Cliinae found during Tara Oceans mission. It is circumglobal. *C. p. pyramidata* is in the border repartition of *C. p. lanceolata.* In the Southern Atlantic Ocean the *pyramidata* group displays high variations: *C. p. sulcata; C. p. martensii.* Some taxa were raised to the species level: *C. convexa, C. antarctica; C. excisa.*

*Hyalaea cuspidata is* present but not abundant in all the oceans (11,4% Cliinae). It was found during the Tara Oceans mission in the Eastern Mediterranean Sea, Northern Indian Ocean, Atlantic Ocean (Eastern South Atlantic, Central South Atlantic, Western South Atlantic, and Western North Atlantic).

C. *recurva* and *C. chaptalii* have a discontinuous global distribution. The bathypelagic species *Clio polita* is caracteristic of the Atlantic Ocean.

#### NOMENCLATURE

*Clio polita* Pelseneer, 1888 was alternately named *falcata* and *polita.*

*Cleodora falcata* Pfeffer, 1880: 96 (priority page) *[Clio falcatum* Meisenheimer 1905b: 422; *Clio falcata* Meisenheimer 1906: 107; *Clio (Euclio) falcata* Bonnevie, 1013: 20)].

*Clio polita* Pelseneer, 1888: 60 *[Clio polita* Meisenheimer 1905a: 20; *Clio (Balantium) polita* Johnson 1934:152; *Euclio polita* Tesch 1946: 15; *Clio polita* Spoel 1967: 75].

According to Tesch (1946) ‘ Meisenheimer (1905b) objected to the use of the name of *polita* on account of it being only a manuscrit designation, so that the name *falcata* Pfeffer was restored by them'. However since Johnson (1934) is generaly choice *polita*.

> Superfamily CAVOLINIOIDEA Gray, 1850
>
> Family CAVOLINIIDAE Gray, 1850
>
> Subfamily **DIACRIINAE** n. subfam.
>
> (Figs 2; 5; 8)

DIAGNOSIS: more or less globular teloconch with lateral ridges and slits, peristome with a thickened short dorsal lip, rudimentary closing system, long juvenile shell (persistant or deciduous) with flat lateral ridges; semi-circular posterior footlobe.

Genera: *Diacria* Gray, 1847 and ***Telodiacria*** n. gen.

They are known to be present from the Miocene (Tortonian) - Pliocene.

Genus: *Diacria* Gray, 1847

Type species: *Hyalaea trispinosa* de Blainville, 1821: 82

*Hyalea trispinosa* Rang, 1829: 115

*Diacria trispinosa* Gray, 1847: 203

*Cleodora infundibulum* Wood, 1842: 459

*Hyaloea trispinosa* Verany, 1853: 380

*Cavolinia trispinosa* Pelseneer, 1866: 346

*Cleodora compressa* Adams, 1853: 52

*Cavolinia trispinosa* (part) Pelseneer, 1866: 346

*Hyalaea (Diacria) trispinosa* Monterosato, 1875: 50

*Pleuropus trispinosa* Pfeffer, 1879: 236

*Cavolinia (Diacria) trispinosa* (part) Dall, 1889: 82

*Diacria trispinosa* Meisenheimer, 1905 a: 2

Species: *Diacria trispinosa* (de Blainville, 1821), *D. major* (Boas, 1886), *D. rampali* Dupont, 1979, *D. maculata* Beeker & Spoel, 1988, *D. piccola* Bleeker and Spoel, 1988, *D. gracilis* Rampal, 2002.

Genus: *Telodiacria* n. gen.

Type species: *Hyalaea quadridentata* de Blainville, 1821: 81

*Hyalea quadridentata* Deshayes & Edwards, 1836: 41

*Cavolina quadridentata* Gray, 1850: 8

*Hyalea costata* Pfeffer, 1880: 91

*Cleodora pygmaea,* Boas, 1886: 84

*Cavolinia quadridentata* Peck, 1894: 453

*Cavolinia (Diacria) quadridentata* Tesch, 1904:36

*Diacria quadridentata* Meisenheimer, 1905a: 29

Species: *Telodiacria quadridentata* (de Blainville, 1821) n. comb., *T. costata* (Pfeffer, 1879) n. comb., *T. danae* Spoel, 1968 n. comb., *T. erythra* Spoel, 1971 n. comb., *T. schmidti* Spoel, 1971 n. comb.

#### MATERIAL EXAMINED

TARA Oceans mission (2009 – 2012): 58 *Diacria* (mainly young spcimens) (34°93’S, 17°93’E). Caribbean Sea (Yucatan; Virgin Islands): 13 *Diacria.*

TARA Oceans mission (2009 – 2012): 132 *Telodiacria* n. gen (mainly young specimens) (21°47S, 54°27’E; 16°09’S, 42°83’E; 20°94’S, 35°19’W).

Caribbean Sea (Yucatan; Virgin Islands): 31

*Telodiacria* n. gen.

#### CLADISTIC AND MOLECULAR ANALYSES

Cladistic: Diacriinae and Cavoliniinae represent two distinct lineages. *Diacria trispinosa* and *Telodiacria quadridentata* n. comb. form a new clade that is the sister group to the Cliinae.

COI data and data set: Diacriinae and Cavoliniinae are not monophyletic; *Diacria* is the sister group to the Cliinae and *Telodiacria* is the sister group to *Diacria* + Cliinae. 28S data and data set (1.00/100): *Diacria* is the sister group to *Telodiacria* and they form a monophyletic subfamily Diacriinae; 28S mol. data (0,97/): the Diacriinae are the sister group to *Cuvierina andHyalocylis.* Despite the incongruence, these analyses support the new genus *Telodiacria* and justify the new subfamily Diacriinae.

Genus: *Diacria* Gray, 1847

Type species: *Hyalaea trispinosa* de Blainville, 1821:81

DIAGNOSIS: flat teloconch, long lateral spines, persistant juvenile shell, spheric protoconch.

#### DEFINITION

*Diacria* has a long, large and flat shell. It includes a large and flat teloconch (ellipsoidal section) with lateral spines and a persistant juvenile shell: l (total shell) = 7.06 – 11,00 mm ; l (teloconch) = 5.44 – 8.69 mm; w (spines) = 5.97 – 11.08 mm; h (teloconch) = 2.05 – 2.91 mm). The very long and narrow juvenile shell has flat lateral ridges and a spheric protoconch. This shell is hyaline and sometimes more or less brown. The median radular teeth have a large and short base with short denticules and triangular curved cuspide (Fig. 8E, F).

The *trispinosa* group is polytypic and polymorphic. *D. trispinosa* (de Blainville, 1821): *D. t. trispinosa* (de Blainville, 1821), *D. t. atlantica* Dupont, 1979 (l = 9.66 – 11mm), *D. t. heterocolorata* Rampal, 2002 (l = 7.06 mm); D. *major* (Boas, 1886) (l teloconch = 7, 40 - 8, 69 mm) (the juvenile shell is often discarded); *D. rampali* Dupont, 1979 (l = 8.72 – 10.48 mm); *D. maculata* Bleeker and Spoel, 1988; *D. piccola* Bleeker and Spoel, 1988; *D. gracilis* Rampal, 2002 (l = 8.86 -10,10 mm) (Rampal 2002).

#### MOLECULAR ANALYSES

COI and 28S: the *Diacria* lineage is monophyletic.These analyses support the taxonomic rank of three species (there is no sequence available for *D. gracilis).* 28S data (1.00/100): *Diacria trispinosa* is the sister group to *D. rampali* + *D. major.* 28S data set (0,99/96): *D. rampali* is the sister group to *D. trispinosa + D. major* (COI data: 0 sequence).

According to Jennings *et al.* (2010) *D. trispinosa* specimens "from widely separated ocean basins were geneticaly similar (E Atlantic and SE Indian); specimens with "sequence differences were detected between regions for *D. major* (NW Atlantic vs. SE Indian)”.

Genus: ***Telodiacria*** n. gen.

Type species: *Hyalaea quadridentata* de Blainville, 1821: 81.

ETYMOLOGY: fets the last development phase telo.

DIAGNOSIS: globular teloconch, lateral spine absent, deciduous juvenile shell, oval protoconch, apex closed by a septum.

#### DEFINITION

Main features: globular hyaline or brown teloconch, absence of lateral spines, presence of a deciduous juvenile shell with an oval protoconch, apex closed by a septum. There are three or five dorso-longitudinal ribs more or less hilly; the peristome has a thickened dorsal lip and transversal striae; the ventral side have transverse striae. Near the lateral fringes there is a vacuolised cells area (Fig. 2H). *Telodiacria quadridentata* n. comb.: l = 2.5 – 3.0 mm; hyaline shell, tickened peristome sometimes brown border; 5 longitudinal slightly swollen dorsal ribs; less proeminent transversal striae.*T. costata* n. comb.: l = 3.5 – 4.0 mm; solide and ticker shell; hyaline sometimes brown shell; 5 deep long and sharp well developed dorsal ribs; well proeminent transversal striae. *T. danae* n. comb.: l = 1.60 – 1,98 mm; hyaline shell; 3 faintly swollen more or less short ribs; transversal striae. We have no observed *T. erythra* Spoel 1971 n. comb. and *T. schmidti* Spoel 1971.

#### MOLECULAR ANALYSES

Limited results because there is a minority of adult specimens. COI (1.00/94, 1.00/95) *Telodiacria* is monophyletic; one young unspecified specimen is the sister group to *T. quadridentata* n. comb. and to *T. danae* n. comb. 28S data set: *T. danae* n. comb. is the sister group to two youg unspecified specimens.

#### DISTRIBUTION

In the Tara Oceans expedition *Diacria* represents 16.4% of the Cavoliniidae. *D. trispinosa atlantica:* Nord Atlantic (30° - 60°). *D. t. trispinosa:* circumglobal (35°N - 35°S) *(other*taxa were certainly assimilated to this specie). *D. rampali:* circumglobal (20°N - 35°S). *D. gracilis:* Western Pacific Ocean. *D. major* is rarely found in all the oceans.

*Telodiacria* (majority of young specimens) represents 20.6 % of the Cavoliniidae. *T. quadridentata* n. comb. and *T. costata* n. comb.: Indo-Pacific area. *T.* danae n. comb. : circumglobal distribution. *T. erythra* Spoel, 1971: Red Sea and Western Indian Ocean.

*T. schmidti* Spoel 1971n. comb. Easten Pacific Ocean.

> Superfamily CAVOLINIOIDEA Gray, 1850
>
> Family CAVOLINIIDAE Gray, 1850
>
> Subfamily CAVOLINIINAE Gray, 1850
>
> (Figs 3; 5)

DIAGNOSIS: more or less globular teloconch with ridges and slits, long peristome with bent dorsal lip, well developed closing system, caracteristic dorsal ribs, juvenile shell absent, wide posterior foot lobe.

Genera: *Cavolinia* Abildgaard, 1791 and *Diacavolinia* Spoel, 1987.

The *Cavoliniinae* can be found from the Miocene (Langhian) onwards

Genus: *Cavolinia* Abildgaard, 1791: 175.

Type species: *Cavolinia tridentata* (Niebuhr, 1775)

*Anomia tridentata* Niebuhr, 1775: 124

*Cavolinia natans* Abildgaard, 1791: 175

*Hyalaea cornea* Lamarck, 1801: 139

*Haloea tridentata* Roissy, 1805: 73

*Hyalea tridentata* Lamarck, 1816: 13

*Cleodora strangulata* Deshayes, 1823: 203

*Pleuropus pellucidus* Eschscholtz, 1825: 735

*Cavolinia (Orbignyia) inflexa* Adams, 1859: 45

*Cavolinia (Cavolinia) tridentata* Tesch, 1904: 37

*Cavolinia* tridentata Spoel, 1967: 94

Species: *Cavolinia gibbosa* (d'Orbigny, 1836) *C. globulosa* (Gray, 1850), *C. inflexa* (Lesueur, 1813), *C. tridentata* (Niebuhr, 1775), *C. uncinata* (Rang, 1829).

Genus : *Diacavolinia* Spoel, 1987: 78

**Type species**: *Hyalaea longirostris* de Blainville, 1821 : 81

*Cleodora strangulata* Deshayes, 1823 : 204

*Hyalea laevigata* Deshayes & Edwards, 1836: 423

*Cavolina longirostra* Gray, 1850: 8

*Pleuropus laevigata* Adams, 1858: 611

*Cavolinia longirostris* Pelseneer, 1888: 79

*Cavolina (Cavolina) longirostris* Dall & Simpson, 1900: 361

*Cavolinia (Cavolinia) longirostris* Tesch, 1904: 41

*Diacavolinia longirostris* Spoel, 1987: 78

**Species: *Diacavolinia longirostris* (de Blainville, 1821), *D. angulosa* Gray, 1850, *D. strangulata* (Deshayes, 1823), *D. limbata* (d’Orbigny, 1836), *D. flexipes* Spoel, 1971,** D. ***mcgowani* Spoel, 1973. Sixteen species are described by Spoel *et al.* (1993): *D. aspina, D. atlantica, D. bandaensis, D. bicornis, D. constricta, D. deblainvillei, D. deshayesi, D. elegans, D. grayi, D. ovalis, D. pacifica, D. robusta, D. souleyeti, D. striata, D. triangulata, D. vanutrechti.***

#### MATERIAL EXAMINED

TARA Oceans mission (2009 – 2012) : 63 specimens (14°59’N, 69°98’E; 06°O3’N, 73°89’E; 21°47S, 54°27’E; 16°95’S, 53°98’E; 16°09’S, 42°83’E; 34°93’S, 17°93’E).

Western Mediterranean Sea, Caribbean Sea (Yucatan; Virgin Islands): 66 specimens.

#### DEFINITION

The Cavoliniinae displays a more or less globular teloconch with an ellipsoidal transversal section; the peristome have a round and curved dorsal lip (straigth and pointed in *Cavolinia inflexa,* beak-like gutter stucture in *Diacavolinia).There* are longitudinal lateral slits with an important closing mechanism (two mecanisms in *Diacavolinia).* The comma-shaped protoconch is in line with the teloconch without clear demarcation; in *Diacavolinia* there is a discarded protoconch and the apex is closed by an irregular slit (no septum) (Spoel *et al.* 1993). *Cavolinia. inflexa* is lower larger near the lateral spines and longer posteriorly.The dorsal ribs number is generaly different in the Atlantic and Indo-Pacific area: *inflexa* group ( 3 or 1 ribs), *gibbosa* group (7 or 5 ribs), *uncinata* group ( 3 or 5 ribs). The parapodia are very elongated and the posterior footlobe is short but very large. The anterior appendages of the mantle are similar to the other Cavoliniidae; the *Diacavolinia* species and *Cavolinia globulosa* also have two antero-ventral trilobed expansions sometimes deciduous (Fig. 3C, E). There are long and retractile lateral appendages (Fig. 4). The pallial gland is similar to the other Cavoliniidae (Fig. 5H). The median teeth of the radula have a triangular base with long denticules and a triangular curved cuspide (Figs 9; 10). *Diacavolinia* species have separate genital aperture (Meisenheimer, 1905).

#### CLADISTIC AND MOLECULAR ANALYSES

In these analyses was named *Diacavolinia* or *Diacavolinia longirostris* seven unspecified specimens (Corse *et al.* 2013).

Cladistic: *Cavolinia* Gray, 1850 and *Diacavolinia* Spoel, 1987 *form* a monophyletic group: Cavoliniinae Gray, 1850.

COI data and data set: the Cavoliniinae are not monophyletic because *Cavolinia inflexa* is basal to a clade composed of the other straight shell Euthecosomata (not including *Hyalocylis).* COI data (0,97/, 0,98/): *Diacavolinia* (Mozambique Channel) is nestled within *Cavolinia labiata* (d’Orbigny, 1836) and within *Cavolinia* sp. (South-Western Indian Ocean). 28S mol. data and gene data set (Figs 15, 16): the Cavoliniinae are monophyletic; these molecular analyses also show affinities of *Diacavolinia* with some *Cavolinia* (1.00/98, 1.00/90): *Diacavolinia* (Carribean Sea) is the sister group to *Cavolinia globulosa* (Gray, 1850) (North Indian Ocean); this group forms a clade monophyletic with the other Cavoliniinae.

#### REMARKS

*Diacavolinia* was validated by DNA barcoding (Mass *et al.* 2013); indeed it has some distinct caracteristics (deciduous protoconch, two closing mechanisms, and distinct genital orifices). Nevertheless the molecular analyses show affinities *Diacavolinia* with *Cavolinia labiata, C. globulosa* and *Cavolinia sp.:* the genus *Diacavolinia* nestled within these *Cavolinia* seems invalidated. Nevertheless it is difficult to explain the different topologies in this subfamily (monophyly or polyphyly), as well as the aberrant position of species to the same group *(Cavolinia inflexa* and *C. labiata)* and the position of *Cavolinia inflexa* (sister group of the entire straight shaped Euthecosomata (except *Hyalocylis).* Further analyses are needed to help understanding these issues. The genus *Diacavolinia* is therefore at present maintained.

#### DISTRIBUTION

In the TARA Oceans samples the Cavoliniinae represents 18% of the Cavoliniidae.

*Gibbosa* group

*Cavolinia gibbosa* (d’Orbigny, 1836): South-Western Indian Ocean, Mozambic Channel and Benguela Current (South-Eastern Atlantic Ocean). *Cavolinia flava* (d’Orbigny, 1836): all over Atlantic Ocean and two rare particular situations: an exceptional incursion into the Western Mediterranean Sea near Gibraltar (Atlantic current indicator) and few presence in the NorthEastern Pacific near Panama: pre-isthmatic Atlantic palaeo-indicator; this last area represents a faunistic entity in the Pacific Ocean (Rampal 2002; 2014). *Cavolinia plana* (Meisenheimer, 1905) is a characteritic Indo-Pacific species. It is found in the Indo-Austral Archipelago (Tesch 1904), troughout the Indo-Western Pacific (Rampal 1975) and only rarely in the South-Eastern Pacific (Meisenheimer 1905). *C. gibboides* Rampal, 2002 is located in the Eastern Mediterranean Sea - South-Tyrrhenian Sea (Rampal 1970b). Unfortunally our molecular analyses did not yield any result for *C. gibboides* (Corse *et al.* 2013). .

*Inflexa* group

*Cavolinia inflexa imitans* (Pfeffer, 1880): cicumglobal distribution. *Cavolinia inflexa inflexa* (Lesueur, 1813): border of the *C. i. imitans* distribution. *Cavolinia labiata* (d’Orbigny, 1836): Indo-Pacific distribution, Mozambique Channel with incursion in the South Eastern Atlantic Ocean. The sporadic presence of *Cavolinia inflexa* in the North-Eastern Pacific Ocean can to touch on the pre-isthmatic palaeo-indicator (Rampal 2002). *Cavolinia labiata* and *C. inflexa* (1.00/99) are found in a same station without hybridization.

*Tridentata* group

*Cavolinia tridentata* (Forskal, 1775) has a special biogeography. It is composed of at least by nine infraspecific members: *tridentata, bermudensis, dakarensis, atlantica, kraussi, danae, teschi, affinis, occidentalis* (Pfeffer 1880; Boas 1886; Schiemenz 1906; Mac Gowan 1960; Spoel 1974; Rampal 1975). Despite a vast polymorphism, two morphological tendencies can be observed. One linked to *tridentata* (large worldwide repartition) and one to *affinis* (d’Orbigny, 1836) (almost localised to the Eastern Pacific Ocean). However, a projection of correspondence analyses shows a succession of affinities between the populations of different oceans. This results in a circumglobal cline with a belt of hybrid populations in the South-Eastern Pacific Ocean (Rampal 1975). Each area has a characteristic phenotype but the morphotype usually shows affinities with the ones from neighbouring areas as a clinal variation.

*Uncinata* group

This tropical group is present in the entire ocean (Mediterranean Sea?). There are Indo-pacific and Atlantic entities. Spoel (1969) described three taxa: *Cavolinia uncinata uncinata* (Rang, 1828) (l = 6.97 mm) and *C. u. roperi* Spoel, 1969 (l = 4.48 mm) from the Atlantic Ocean; *C .u. pulsata* Spoel, 1969 (l = 8.14 mm) from the Indian Ocean. These taxa differ by the dorsal ornementation: The Atlantic specimens have three dorsal ribs (the two laterals very lightly deprimed on the middle); the indo-pacific specimens have five dorsal ribs; the Indian specimens are the biggest (Rampal 1979). According to Jennings *et* al.(2010) in COI there is no subtantial regional variation among Atlantic locations for *C. uncinata* but sequence differences were detected between different regions "(Atlantic vs.SE Indian)”.

*Longirostris* group

The polymorphic and polytypic recent genus *Diacavolinia* is present in all the oceans (rare in the Mediterranean Sea). According to Spoel *et al.* (1993) there are characteristic repartitions: Atlantic Ocean: *D. atlantica, D. deblainvillei, D. ovalis, D. deshayesi, D. constricta, D. strangulata, D. longirostris, D. limbata.*

Indian Ocean: *D. bicornis, D. souleyeti, D. striata, D. aspina.*

Red Sea: *D. flexipes.*

Pacific Ocean: *D. pacifica, D. vanutrechti, D. triangulata, D.* robusta, *D. mcgowani,* Malaisie: *D. vanutrechti*, *D. elegans*, *D. bandaensis*.

Atlantic, Indian and Pacific Ocean: *D. angulos.*

*Cavolinia globulosa* (Gray, 1850): Indo-Pacific species.

> NEW CLASSIFICATION (extant taxa)

THECOSOMATA de Blainville, 1824

EUTHECOSOMATA Meisenheimer, 1905

Family Limacinidae Gray, 1847

Genus *Limacina* Bosc, 1817;

*Limacina helicina* (Phipps, 1774)

Family Heliconoididae n. fam.

Genus *Heliconoides* d'Orbigny, 1835; *Heliconoides inflata* (d'Orbigny, 1835)

Family Thieleidae n. fam.

Genus *Thielea* Strebel, 1908; *Thielea helicoides* (Jeffreys, 1877)

Superfamily CAVOLINIOIDEA Gray, 1850

Family Creseidae Rampal, 1973

Genus *Creseis* Rang, 1828; *Creseis virgula* (Rang, 1828)

Genus *Boasia* Dall, 1889; *Boasia chierchiae* (Boas, 1886) n. comb.

Family Cavoliniidae Gray, 1850

Subfamily Cuvierininae Gray, 1847

Genus *Cuvierina* Boas, 1886; *Cuvierina columnella* (Rang, 1827)

Subfamily Cliinae Jeffreys, 1869

Genus *Clio* Linnaeus, 1767; *Clio pyramidata* Linnaeus, 1767

Genus *Hyalaea* de Blainville, 1821 ; *Hyalaea cuspidata* Bosc, 1802 Subfamily Diacriinae n. subfam.

Genus *Diacria* Gray, 1847; *Diacria trispinosa* (de Blainville, 1821)

Genus *Telodiacria* n. gen.; *Telodiacria quadridentata* (de Blainville, 1821) n. comb.

Subfamily Cavoliniinae Gray, 1850

Genus *Cavolinia* Abildgaard, 1791; *Cavolinia tridentata* (Niebuhr, 1775)

Genus *Diacavolinia* Spoel, 1987 ; *Diacavolinia longirostris* (de Blainville, 1821)

Genera *Incertae sedis*

Genus *Styliola* Gray, 1850; *Styliola subula* (Quoy & Gaimard, 1827)

Genus *Hyalocylis* Fol, 1875; *Hyalocylis striata* (Rang, 1828)

### Phylogenetic hypotheses

Based on morphological and anatomical data of the Thecosomata Euthecosomata, two hypotheses for the appearance of the straight shell were proposed: single appearance for the Cavolinioidea Gray, 1850 from a single spiral shell (Boas 1886, Pelseneer 1888, Meisenheimer 1905, Bonnevie 1913) or two independent appearance for the Creseidae Rampal, 1973 and the Cavoliniidae Gray, 1850 from two spiral shell lineages *Limacina* Bosc,1817 and *Thielea* Strebel, 1908 which have a same hypothetical benthic ancestor lineage (Rampal, 1973).

Our recent cladistic and molecular analyses present two hypotheses. The analyses in COI tree support the double emergency: the straight shell species could likely have appeared twice independantly with a reversion in the lineage *Thielea / Heliconoides* to explain the coiled shell observed in these genera (Corse *et al.* 2013).The double emergency also was shown in the COI tree by Jennings *et al.* (2010) (these authors do not offer any phylogenetical conclusion on account of 'their poor taxonomic sampling’). However this scenario was not adopted because it is the less parsimonious: the unwinding of the spiral shell species is homoplasic. It is the less informative for the older divergences. COI showed a lack of phylogenetic signal to infer Euthecosomata relationships (similar problem for other Mollusca).Therefore this hypthesis of the double emergency was declined (Corse *et* al.2013).

The analyses in 28S / morphological data also propose the monophyly of the Cavolinioidea Gray, 1850: the straight shell species originated from a unique coiled shell ancestor that subsequently evolved into a straight conical morphotype belonging to the genus *Creseis* ancestral to the more complex straight shells of the Cavoliniidae Gray, 1850. This hypothesis is the most parsimonious analysis: it includes the unwinding in one time, it is the most informative for older divergence and it is more reliable and congruente with palaeontological data for revolving the deep nodes. We have kept this hypothesis of the monophyly (Corse *et al.* 2013). Nevertheless the incongruence between gene trees could also be the result of incomplete data for reconstructing phylogenies. Therefore these conclusions require some reserves and needs further investigations (Corse *et al.* 2013).

The spiral shell species notably, were considered neotenic organisms derived from spiral benthic ancestors (Lemche 1948; Huber 1993), hypothesis not fully accepted notably by Lalli and Gilmer (1989).

The transition between the spiral shell forms and the straight ones is illustrated by the existence of partially unwounding fossils *Camptoceratops* Wenz, 1923 and *Sphaerocina* Jung, 1971. This transition performed by a 180° twist of the trunk relative to the head, and an unwinding of the visceral mass (Boas 1886) was observed to some recent species during the ontogenesis (Fol 1875). The Euthecosomata with spiral shell are known to be present from Upper Paleocene-Lower Eocene (Watelet & Lefèvre 1885). The first with straight shell is conical that is still in the first process unwinding spiral shell; it emerged during the Lower Eocene (Hodgkinson *et al.* 1992; Curry 1965). The hypothesis for the impact of this macro-unwinding of the spiral shell (which have prismatic and crossed lamellar aragonitic fibres) over the microstructural aragonitic straight shell can be seen in their spiral aragonitic microstructure (Rampal 1975). This spiral microstructure is already present in the partially unwound fossil *Camptoceratops priscum* (Gowin-Austen, 1925) (Curry & Rampal 1979) and signs of it can be observed in the apex protoconch of *Heliconoides inflata* (Glaçon *et al.* 1994, pl. 3, Figs 1-9) and in the apex of *Limacina retroversa* (Fig. 6F); this spiral microscruture in the protochonch seems to anticipate the macro-unwinding. In the meroplanktonic veligers of some benthic Tectibranch molluscs it is observed the same aragonitic structure (Richter 1976). According to Haszprunar (1985) the structure of the Euthecosomata illustrates the paedomorphic aspect of the shell's long development.

## DISCUSSION

The emergence of the holoplanktonic Euthecosomata from benthic Gastropoda occurred through successive radiations from the Upper Paleocene – Lower Eocene until the Pliocene. These events involved deep and important morphological and anatomical changes: the acquisition of swimming organs in the spiral shell Euthecosomata and then transition to a straight shell, which later was gradually optimised. Some taxa became extinct or their diversity decreased during the Tertiary era, however, interestingly, *Limacina* Bosc, 1817, *Creseis* Rang, 1828 and *Cuvierina* Boas, 1886 are always present amongst the earliest genera (Eocene). The present study gave an account of these diversifying events. Moreover, a discriminating between phylogenetic hypotheses and a new systematic of the Euthecosomata was submitted. Two phylogenetic hypotheses were originally considered. The first one proposes a double emergence of straight shell species from two separate lineages of spiral shell ancestors (Rampal 1973). According COI tree there is the double emergency. However 28S tree did not support this scenario wich is the less parsimonius; the second hypothesis we have kept *assumes* the monophyly of Euthecosomata (Corse *et al.* 2013). The congruence of our results from cladistic, molecular (28S) and palaeontological analyses supported this interpretation: a first transition from a spiral shell morphotype into the first plesiomorphe straight conica shell *(Creseis)* wich in turn is ancestral to all the other genus with apomorphic straight shells. It should be emphasised that future studies of currently non available 28S sequences for *Heliconoides inflata* (d’Orbigny, 1835) and *Thielea helicoides* (Jeffreys, 1877) may impact on these interpretations, require some reserve and need further investigations.

New classification. Spiral shell Euthecosomata: Limacinidae Gray, 1847; Heliconoididae n. fam.; Thieleidae n. fam. Straight shell Euthecosomata: Cavolinioidea Gray, 1850: Creseidae Rampal, 1973; Cavoliniidae Gray, 1850: Cuvierininae Gray, 1847; Cliinae Jeffreys, 1869; Diacriinae n. subfam.; Cavoliniinae Gray, 1850. The genera *Styliola* Gray, 1850 and *Hyalocylis* Fol, 1875 are considered *genus incertae sedis.*

New genera are proposed or reinstated: *Telodiacria* n. gen., *Hyalaea* Bosc, 1802 and Boasia Dall, 1889.

Most of the species have their taxonomic rank confirmed. The fossil *Altaspiratella* Korobkov, 1966 was excluded from the Limacinidae while the fossil *Vaginella* Daudin, 1800 was integrated with the Cuvierininae. Two diversifying events of interest should be highlighted: the late appearance of *Hyalocylis* (Upper Miocene) despite its lineage starting earlier (Lower Oligocene) and the neoteny process illustrated by *Styliola.* The integrative approach to the estimation of divergence times, which explained the incongruences concerning *Hyalocylis* is a powerful tool and could help elucidate other issues, such as the late emergence of archaic phenotypical species *Thielea helicoides* (Jeffreys, 1877) and *Clio polita* (Pelseneer, 1888).

## ABBREVATIONS

h: height
l: length
t: thickness
w: width
MA: millions years
ICZN: INTERNATIONAL COMMISSION ON ZOOLOGICAL NOMENCLATURE

## ACKNOWLEDGMENTS

I wish to thank Emmanuel Corse for the molecular analyses and his participation to their interpretation with André Gilles, Yvan Perez and Nicolas Pech (Evolution Génome Environnement, Aix-Marseille Université, France).I am grateful to Philippe Bouchet and Pierre Lozouet (Museum national d'Histoire naturelle, Paris) for the advice on the zoological terminology. I am grateful to the reviewers for the fundamental comments and criticism. I wish to thank Daniel Papillon for the English translation. I also thank Fabien Francl for processing the iconography.

